# Planetary microbiome structure and generalist-driven gene flow across disparate habitats

**DOI:** 10.1101/2025.07.18.664989

**Authors:** Chan Yeong Kim, Daniel Podlesny, Jonas Schiller, Supriya Khedkar, Anthony Fullam, Askarbek Orakov, Christian Schudoma, Shahriyar Mahdi Robbani, Anastasiia Grekova, Michael Kuhn, Peer Bork

**Author notes:** Corresponding author: Peer Bork. These authors contributed equally to this work (Chan Yeong Kim, Daniel Podlesny, Jonas Schiller).

## Abstract

Microbes are ubiquitous on Earth, forming microbiomes that sustain macroscopic life and biogeochemical cycles. Microbial dispersion, driven by natural processes and human activities, interconnects microbiomes across habitats, yet most comparative studies focused on specific ecosystems. To study planetary microbiome structure, function, and inter-habitat interactions, we systematically integrated 85,604 public metagenomes spanning diverse habitats worldwide. Using species-based unsupervised clustering and parameter modeling, we delineated 40 habitat clusters and quantified their ecological similarity. Our framework identified key drivers shaping microbiome structure, such as ocean temperature and host lifestyle. Regardless of biogeography, microbiomes were structured primarily by host-associated or environmental conditions, also reflected in genomic and functional traits inferred from 2,065,975 genomes. Generalists emerged as vehicles thriving and facilitating gene flow across ecologically disparate habitat types, illustrated by generalist-mediated horizontal transfer of an antibiotic resistance island across human gut and wastewater, further dispersing to other habitats, exemplifying human impact on the planetary microbiome.

## Introduction

Microbes are essential members of almost every ecosystem on our planet. They usually form communities, microbiomes, that drive ecosystem processes, biogeochemical cycles^1^, and sustain macroscopic life, often through direct associations with their hosts^2^. Although most microbial species and genes are habitat-specific^3–5^, various mechanisms facilitate their dispersal at different temporal and spatial scales, interconnecting microbiomes across diverse and geographically distant habitats^6–9^. Humans amplify this connectivity, exemplified by the rapid spread of infections and antibiotic resistance genes across the globe^10–12^. Microbial dispersal and environmental selection fundamentally shape microbiome structure (taxonomic composition)^13–15^, which is composed of an evolving set of lineages that form a gradient between strictly habitat-specific species (specialists) and species that thrive in multiple and disparate habitats (generalists). However, the extent to which species traverse across habitats and the mechanisms by which they mediate gene flow at a planetary scale remain unclear.

Numerous large-scale studies of microbiome structure and function have been conducted, yet most focus on specific ecosystems (e.g., US riverine^16^, global ocean^17^, topsoil^18^, and human intestine^19^), are restricted to a national scale^20^, are relatively sparse^21^, or are limited in resolving fine-grained taxonomy^19,22^. Moreover, habitat annotations and contextual information are often incomplete or inconsistent, and human-defined ontologies of environments^23,24^ may not always align with microbial habitats. Recently, databases such as SPIRE^24^ or MicrobeAtlas^25^ have emerged, offering uniformly processed sequencing data and annotations to facilitate planetary-scale comparative microbiome analyses. However, to fully leverage such resources, a standardized planetary-scale framework is necessary to contextualize habitat types and quantify their connectivity, disentangle the local, regional, and global drivers influencing the transmission of species and flow of genes, and better understand the evolution of microbiome structure (specifically at species level) and function (specifically at gene level).

In this study, we employed an unsupervised species-based clustering and parameter modeling approach on 85,604 samples from the SPIRE database, which is the largest integration and systematic analysis of shotgun metagenomic data to date. We identified 40 microbial habitat types (hereafter referred to as habitat clusters) that broadly align with existing environmental ontologies. We quantified their ecological similarity and provided a comprehensive hierarchical annotation that reflects associated functional traits and known environmental conditions. This framework enables the study of fundamental questions in microbial ecology^26^, e.g., how to trace microbial dispersal, how microbiomes are shaped, and how they are affected by human activities.

We integrated data from each of the 40 habitat clusters to compare their structural and functional composition. To systematically study gene flow on a planetary scale, we captured both horizontal gene transfer between sympatric species, as well as species transmission across habitat clusters. By devising a robust scoring scheme to quantify the gradient from prokaryotic specialism to generalism, we identified generalists as critical vehicles for gene flow between ecologically disparate habitat clusters. This is illustrated by the identification of an antibiotic resistance island, widely shared among human gut specialists and a generalist species, that was able to spread across gut, wastewater, lake, and estuary microbiomes. Collectively, our findings highlight the profound ecological and health implications of microbial gene flow, to which humans contribute.

## Results

### Prokaryotic species composition reveals habitats at high resolution

To delineate microbial habitats and associated functional traits, we adopted a data-driven metagenomic approach. Leveraging the SPIRE resource^24^, we utilized a set of 85,604 quality-filtered metagenomic samples, collected across the globe. To identify metagenomic features and parameter conditions under which samples would naturally group into ecologically meaningful habitats, we systematically tested an extensive array of input profiles (taxonomic, functional, k-mer-based), dimensionality reduction techniques and unsupervised clustering methods (**Figure 1A**, **Methods**, **Supplementary Table 1**).

**Figure 1.**
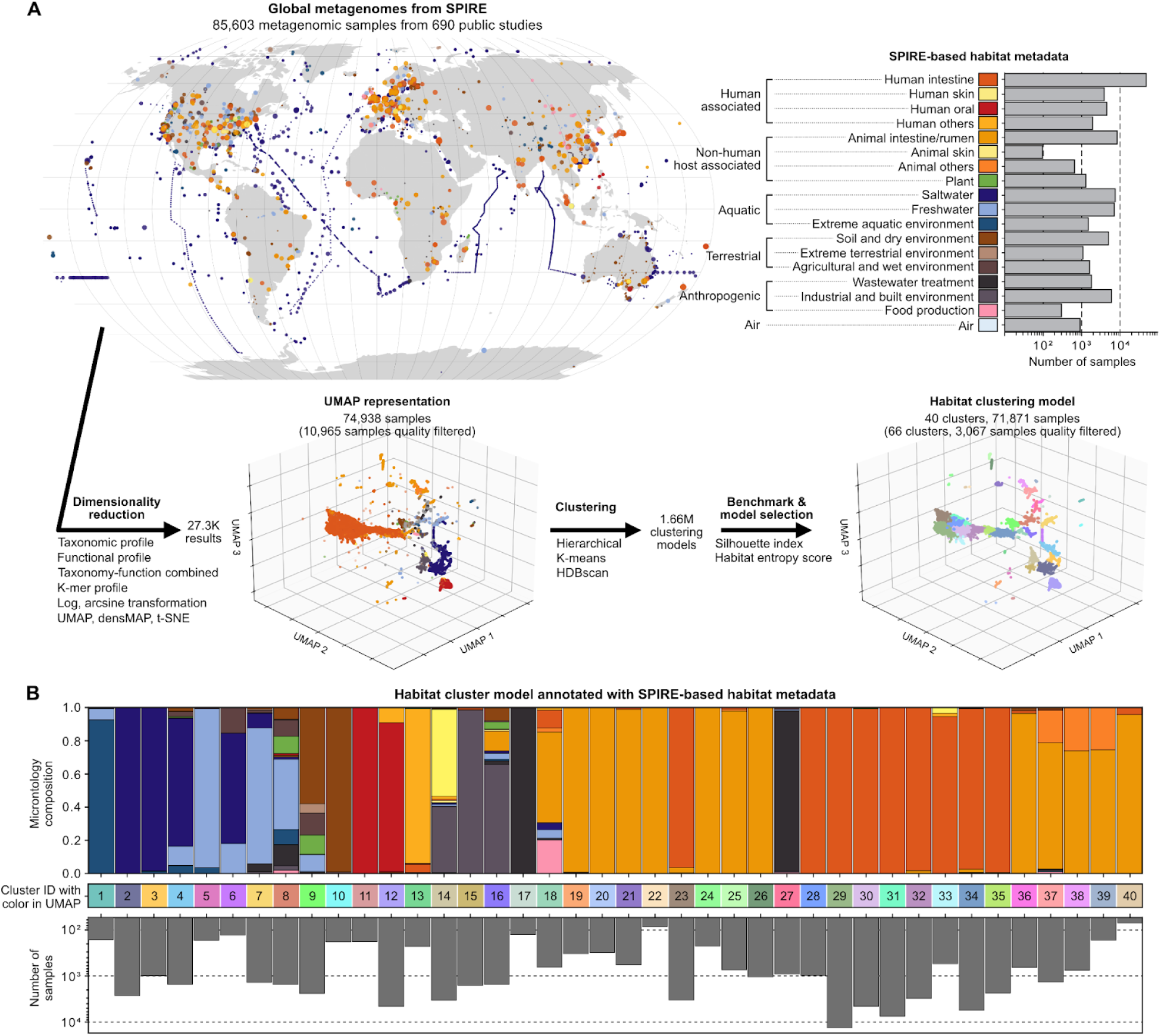
Planetary-scale framework for microbiome-based habitat clustering. **(A)** Overview of the sample collection, dimensionality reduction, clustering, and benchmarking processes. The upper panels show the geographical distribution of the metagenomic samples analyzed in this study and the statistics of simplified microntology categories. 3D UMAP embedding based on the inter-sample Bray-Curtis dissimilarity of species compositions, used for clustering and downstream analyses (**Methods**). **(B)** Visualization of the 40 clusters emerging from the best-performing habitat cluster model, including the sample count per cluster, cluster IDs from the UMAP embedding (A), and composition according to the SPIRE-based habitat microntology (A). Since the habitat entropy score penalizes fusion but not fission of broader habitat terms, these were split into multiple clusters (e.g., human and animal intestine/rumen) or fused when their taxonomic compositions were similar (e.g., cluster 14 composed mostly of human skin and surface swab samples).

We evaluated more than 1.66 million models using metrics that assess both cluster consistency and homogeneity, based on broad and high-level manually curated habitat annotations (**Methods**). Because marked differences in microbiomes among broad habitats (e.g., ocean, terrestrial, skin, and gut) are evident^5,22,27^, we imposed a penalty on the fusion of broad habitat categories. However, to allow for detection of substructures within broad categories, we did not penalize their fission (**Methods**). This comprehensive parameter exploration revealed that applying UMAP^28^ (Uniform Manifold Approximation and Projection) to Bray-Curtis distances between species-level taxonomic profiles most effectively captured subtle differences among habitats (**Supplementary Figure 1A, B**). This effect was further enhanced by logarithmic and to a lesser extent by arcsine scaling, both of which emphasize lower-abundance microbes within taxonomic profiles (**Supplementary Figure 1C, D**). While previous research reported that functional profiles provide lower resolution than taxonomic profiles in human microbiomes^29^, we show that this pattern extends across microbiomes from a wide range of environments: Functional profiles, at the resolution of Clusters of Orthologous groups (COG), enabled broad distinctions such as host-associated versus environmental samples, but failed to resolve finer-scale differences (**Supplementary Figure 2A, B**). Similarly, k-mer-based approaches^30^, though more discriminative than functional profiles, did not surpass the specificity achieved by taxonomic data (**Supplementary Figure 2C, D**). Based on the parameter exploration, we selected the best-performing model, which we further curated to eliminate artifacts arising from individual studies or methodological biases (**Methods**, **Supplementary Figure 3**). The remaining 71,871 metagenomic samples formed 40 robust habitat clusters (**Figure 1B**, **Supplementary Figure 4**, **Supplementary Table 2**).

### Host and environmental filtering structures global microbial communities

As species are mostly habitat-specific^4,5^, each of the 40 clusters likely reflects a distinctive habitat condition. To construct a planetary-scale map of microbial habitats, we evaluated their ecological distances and annotated the clusters using fine-grained, manually curated contextual data where available (**Figure 2A**). Since UMAP prioritizes preserving local over global structure, inter-cluster distances in UMAP embedding do not reflect true ecological distances. Therefore, we used the overlap coefficient of microbial taxa to quantify inter-cluster ecological distances (**Methods**). This revealed a primary structuring of all studied microbiomes into host-associated (**Figure 2A**-clade m) and environmental (**Figure 2A**-clade d) groups, regardless of the geographic sampling locations. Within these primary structures, we identified further fine-grained differentiation shaped by large-scale environmental gradients or host-specific conditions, each of which left distinct signatures in the microbiome structure. We were able to contextualize these signatures with previous literature, indicating the robustness of the framework, as illustrated with three examples: (i) human gut habitat clusters, reflecting age and lifestyle^31^; (ii) animal gut habitat clusters, shaped by host and their diet^32,33^; and (iii) marine habitat clusters, primarily structured by temperature rather than geographic proximity^17^ (**Figure 2**, cf. **Supplementary Information** on major environmental factors for other habitat clusters).

i. Since the human gut accounted for the largest number of samples (46,971), we achieved particularly high resolution in delineating its microbiome structure. We identified nine distinct human gut habitat clusters that separated along known age-related and lifestyle gradients. The adult fecal microbiomes were represented in human gut clusters HG1–HG5 and HG7 (for nomenclature see **Figure 2A**) and clearly separated from a clade with newborn gut (NBG) and infant gut (IFG) species profiles (average age 15 days and 7 months, respectively; **Figure 2A**-clade i). Following the age trajectory, the HG6 cluster (average age of 2 years) was already more similar to the adult microbiomes^34–36^ (**Figure 2B, C**). In adulthood, clusters no longer segregated by age, but lifestyle differences became increasingly prominent. Notably, HG1 was distinctly segregated from other human gut habitat clusters (**Figure 2A, B**), comprising fecal samples predominantly from hunter-gatherers, indigenous Malaysian populations, paleofeces, and lower-income countries^37^, thus distinguishing it socioeconomically and culturally from other gut microbiomes (**Supplementary Figure 5A, B**). This aligns with reports that individuals with traditional lifestyles tend to have a markedly different gut microbiota composition from those in industrialized settings^38^. Enrichment of samples from specific geographical regions was also evident in HG2, HG3, and HG5, which were dominated by samples from Israel, Europe, and East Asia, respectively (**Supplementary Figure 5A**). Thus, the planetary-scale microbiome clustering captured the major conditions underlying human gut microbiome variation.
ii. Animal gut clusters exhibited a high degree of host specificity (**Figure 2A**), each comprising samples from a single animal genus or closely related genera, with the exception of a heterogeneous animal cluster (ANI) (**Figure 2D**). Primate (MKG for monkey, APG for apes) and pig (PG1, PG2) clusters were taxonomically closer to the human gut clusters, whereas herbivores (BG1–BG3 for bovines and HRG for horses) and rodents (MOG for mouse and RMG for rat and myodes) formed distinct clades distant from the human gut microbiota (**Figure 2A**). These observations substantiate previous findings that host diet is a stronger driver of microbial community composition than host evolutionary distance^32,33^.
iii. Marine samples grouped into three habitat clusters (OHL, OIL and OLL representing high, intermediate, and low absolute latitude, respectively) with significantly different absolute latitude ranges (two-sided Mann-Whitney U test *p* < 1e-90) (**Figure 2E–G**). Samples from temperate to polar zones from both hemispheres clustered together, reflecting similarities in species-level taxonomic profiles despite vast geographic distances^39^. To assess the influence of environmental factors (e.g., temperature, depth, pH, salinity, dissolved oxygen, and others) on the ocean microbiome structure, we compared them across clusters using a Kruskal-Wallis test. Temperature had the highest H-statistic value (1571.77, see also **Figure 2H**), more than three times higher than the next most distinguishing factors, salinity (424.01) and dissolved oxygen (413.15) (**Figure 2I**), confirming temperature as the primary factor shaping global marine microbiome structure^17^.

**Figure 2.**
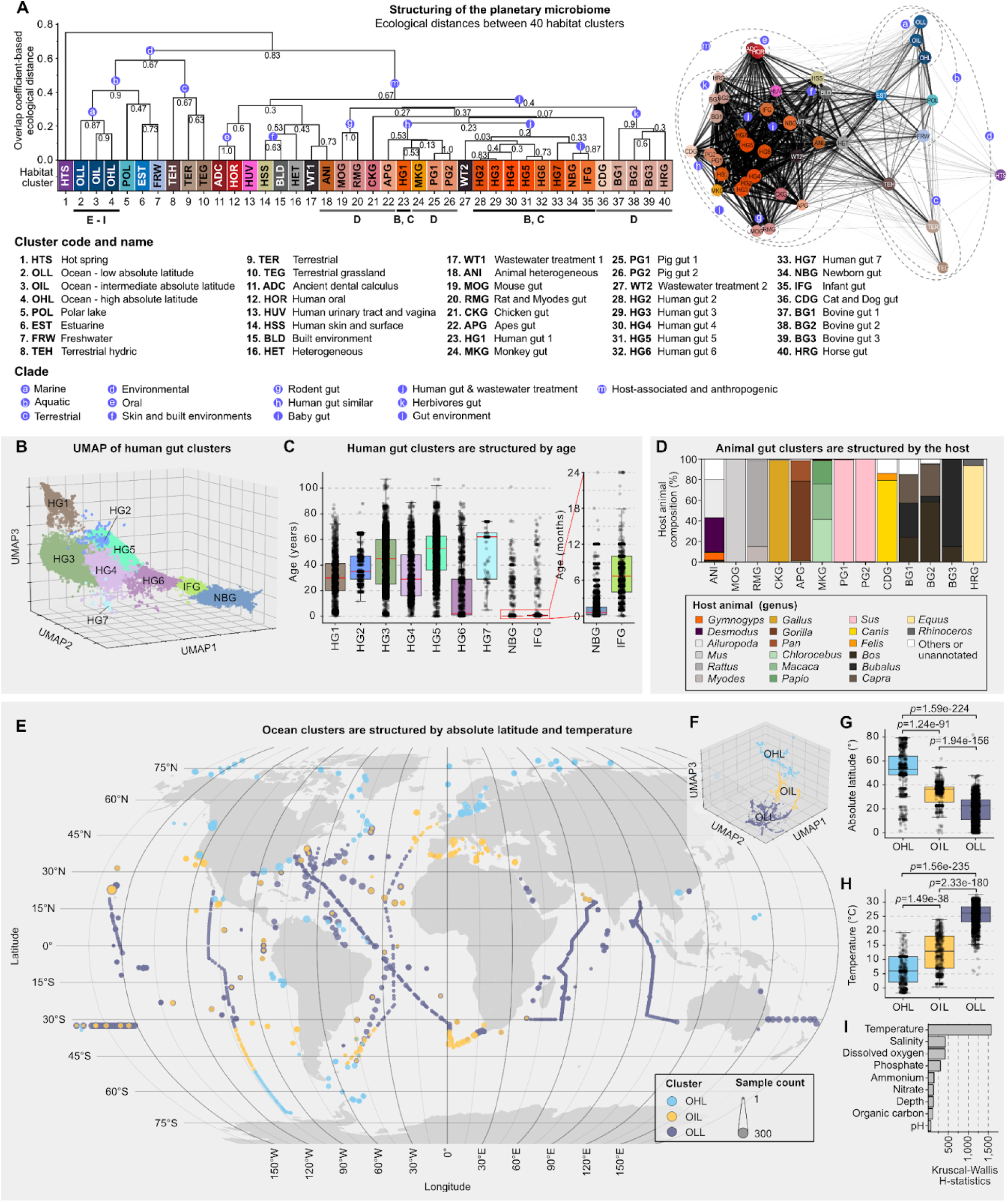
Annotation, ecological distances, and sample characteristics of the habitat clusters. **(A)** Primary structuring of the planetary microbiome into host-associated and environmental habitats, interconnected by anthropogenic microbiomes. Dendrogram (left) and network (right) visualization showing the Ecological distances between habitat clusters were determined by the (dis)similarity of taxonomic profiles. Dendrogram: Hierarchical clustering of the 40 habitat clusters, quantifying ecological distance based on the overlap coefficient of microbial taxa (**Methods**). Values at the tree nodes represent how consistently each clade is preserved across different prevalence cutoffs. Network: Nodes represent the habitat clusters and edges represent the ecological distances between them. The positions of the nodes were determined by Prefuse Force-Directed Layout using negative logarithmic ecological distance as weights (**Methods**). Clusters were annotated based on the characteristics of their constituent samples (inferred from fine-grained contextual data) and are represented by three-letter abbreviations. Letters (B–H) below the cluster indices link to downstream panels. **(B–C)** Human gut clusters are structured by age: (B) UMAP embedding of the nine human gut clusters shows an age-dependent gradient from newborns (NBG) to infants (IFG) and adults (HG1–HG7), (C) Age distribution of hosts from which human gut cluster samples were collected. The box plot on the right zooms into the age distribution for the newborn gut (NBG) and infant gut (IFG) clusters. 91.22% of the NBG samples were annotated as being from individuals aged 6 months or younger, and 93.63% of the IFG samples were from individuals aged 24 months or younger. **(D)** Animal gut clusters are structured by the host animal. For example, the monkey gut cluster (MKG) consists of microbiome samples from *Chlorocebus*, *Macaca*, and *Papio*, which are all members of the *Cercopithecidae* family. Composition of the host animal phylogeny (at the genus level) in non-human animal gut clusters. **(E–I)** Ocean samples cluster by absolute latitude: (E) Geographical distribution, (F) UMAP embedding, (G) absolute latitude, and (H) temperature of samples in three ocean clusters (OHL, OIL, and OLL), (I) Kruskal-Wallis H-statistics for environmental variables comparing the three ocean clusters. (G, H) *p*-values were calculated using the two-sided Mann–Whitney U test.

The robustness of our approach enabled us to flag potential contamination and to classify habitat clusters when habitat annotations were missing from the public data. For example, mapping of metagenomic samples lacking host annotations to habitat clusters accurately inferred the host animal, as validated by host DNA contamination analysis (**Methods**, **Supplementary Figure 6A**), and aquatic samples clustered into the human skin cluster (HSS) (**Supplementary Figure 6B**) exhibited significantly higher human DNA content compared to correctly clustered samples (two-sided Mann-Whitney U test *p* < 1e-09, **Supplementary Figure 6C**).

### Habitat conditions select for microbial functions and genomic traits

Metagenomics offers not only high taxonomic resolution but also rich functional information, providing insights into genomic and phenotypic traits of microbes. To examine how these traits shape structural differences between habitat clusters, we analyzed 2,065,075 isolate genomes and high-quality metagenome-assembled genomes (MAGs) from species within those clusters (**Figure 3, Supplementary Figure 7, Methods**).

**Figure 3.**
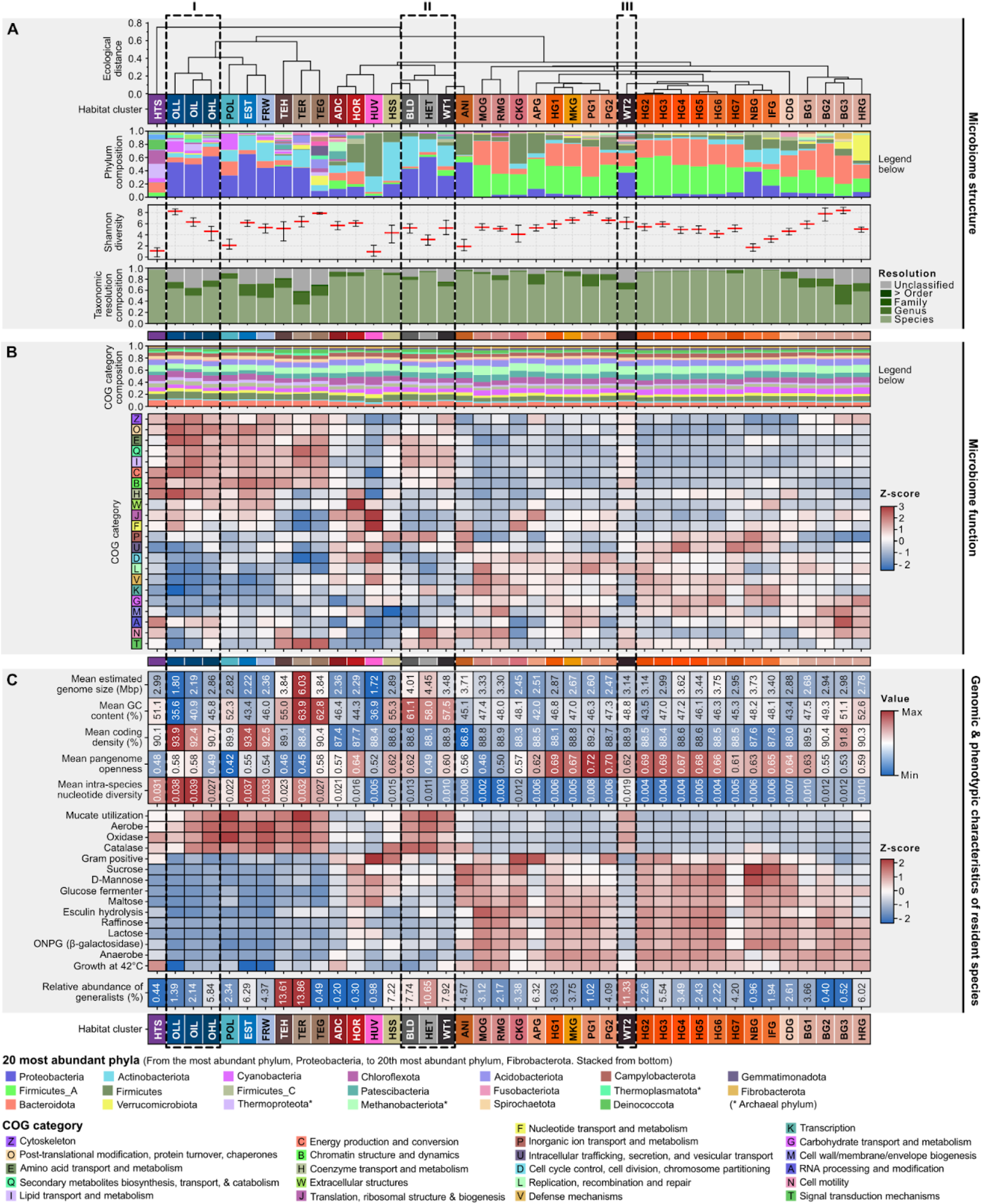
**Microbiome structure, function, and characteristics of species within habitat clusters** The planetary microbiome structure is mirrored in the functional composition of habitat clusters and in the genomic and phenotypic traits of the species residing in them. **(A)** Taxonomic properties of habitat clusters. Dendrogram of ecological distance (see Figure 2A), phylum level composition, Shannon diversity, and taxonomic classification resolution (median with interquartile range), from top to bottom. Host-associated microbiomes were predominantly composed of Firmicutes_A and Bacteroidota, whereas environmental habitats were enriched in Proteobacteria and Actinobacteriota. **(B)** Functional properties of habitat clusters summarized at COG-category level show contrasting patterns in host-associated and environmental microbiomes. Raw COG category composition is shown as a stacked bar plot (top), and z-score normalized per category values are shown as a heatmap (bottom). **(C)** Genomic (upper panel) and phenotypic traits (center panel) of species residing within habitat clusters. Only the top 15 phenotypic traits with the highest variability across the habitat clusters are shown (**Supplementary Figure 7** for the complete set of traits). These traits mirror habitat conditions: For example, frequently oxygenated environments such as aquatic and terrestrial habitats show a high proportion of aerotolerant (Aerobe) microbes. Proportion of generalist species (bottom panel, generalism score ≥ 0.1) per habitat cluster (Figure 4 & **Supplementary Figure 8** for a detailed introduction into the generalism score).

Taxonomic composition differed markedly between habitat clusters, even at the phylum level. Whereas Firmicutes_A and Bacteroidota dominated host-associated microbiomes, Proteobacteria and Actinobacteriota were prevalent in environmental habitats. Rarefaction analysis revealed that at equal numbers of samples, all host-associated habitats (**Figure 2A**-clade m) together (but not individually) harbored greater diversity of species compared to terrestrial (**Figure 2A**-clade c) and aquatic (**Figure 2A**-clade b) microbiomes. While host-associated microbiomes support a high number of closely related species^40,41^ (**Supplementary Figure 8A**), at the genus level, a broader diversity of lineages was observed across aquatic and terrestrial habitats (**Supplementary Figure 8B**). Rarefaction analysis further indicated the ongoing discovery of taxa in terrestrial, aquatic and animal gut microbiomes (but to a lesser extent in the human gut microbiome) (**Supplementary Figure 8A, B**), suggesting numerous undiscovered taxa in these habitat types.

Functional composition, based on broad Clusters of Orthologous Groups (COGs), was more conserved across the habitat clusters than taxonomic composition (**Figure 3A, B**), which might be due to a large fraction of shared core functionalities between microbial communities^17,42^. Still, there were marked differences between host-associated (**Figure 2A**-clade m) and environmental (**Figure 2A**-clade d) habitat clusters (**Figure 3B**). Environmental habitat clusters exhibited higher proportions of cellular processes (COG categories B, O, W) and metabolic functions (COG categories C, E, H, I, Q), pointing to high metabolic versatility (**Figure 3B**). They were also enriched in traits related to aerobic conditions, whereas human and animal gut clusters were dominated by species with anaerobic lifestyles and a higher potential to utilize sugars like sucrose, D-mannose, glucose, maltose, raffinose, and lactose (**Figure 3C**).

Microbes in each habitat optimized their genetic repertoires differently: Environmental microbiomes showed high intra-species nucleotide diversity but closed intra-species gene pools (pangenomes) (**Figure 3C**), indicating more frequent adaptation through point mutations than horizontal gene transfer (HGT). Host-associated microbiomes, conversely, exhibited low intra-species nucleotide diversity and often open pangenomes, especially in the gut of omnivorous animals (humans, pigs, apes, and monkeys) (**Figure 3C**), suggesting that they instead rely more on HGT for functional innovation which requires stronger physical proximity^43^. Genome sizes of host-associated microbiomes were constrained to an upper limit of around 3.5 Mbp, with the human urinary tract and vagina cluster (HUV) displaying the smallest average genome size (1.72 Mbp) (**Figure 3C**). Aquatic clusters had compact genomes (< 2.86 Mbp) with high coding density (90–94%) and low GC content (< 46%), whereas terrestrial clusters featured genomes that are less streamlined with larger genomes (> 3.84 Mbp), lower coding density (88–90%), and high GC content (> 55%) (**Figure 3C**). This is consistent with previous reports^44–46^ and may reflect a trade-off from balancing the energetic costs of genome maintenance with the need for adaptive potential under fluctuating habitat conditions. Since these patterns are visible at the level of habitat clusters, habitat-specific conditions may play a dominant role in shaping these optimization strategies and thereby genomic characteristics^47^.

We observed three ocean clusters (OIL, OLL, and OHL) separated along distinct gradient patterns. The latitude segregation correlated with taxonomic Shannon diversity, peaking in low latitude tropical and subtropical regions (OLL), and declining towards intermediate latitudes beyond the subtropics (OIL) and high latitudes (OHL) (**Figure 3A**, dashed rectangle i). This transition was accompanied by an increase in genome size and GC content of the microbes inhabiting these communities, consistent with previous findings^48^ (**Figure 3B**). Additionally, the proportion of aerobic microbes increased towards high latitudes, coinciding with higher dissolved oxygen levels in colder waters (**Figure 3C**). These results highlight the impact of gradient-dependent environmental filtering on the function and diversity of the ocean microbiome.

Together, host and environmental filtering of the microbiome structure was reflected by functional composition and genomic adaptation processes (e.g., pangenome openness, intra-species nucleotide diversity, genome streamlining). Therefore, microbes in each habitat cluster employ fundamentally different strategies to adapt to their environment, likely influenced by abiotic (e.g., nutrient availability, temperature, oxygen) and biotic (e.g., species interaction, competition) factors.

### Metabolic versatility and oxygen utilization are key features associated with prokaryotic generalism

Species adaptation to the environment is often a gradual process and constant species dispersal creates opportunities for colonization. As metagenomic samples represent only a snapshot of this process, the observed species usually span a wide spectrum, from specialists that have adapted and thrive in specific habitats to generalists capable of tolerating a broad range of environmental conditions. To quantify this continuum, we used our framework capturing microbiome structure to develop a generalism score that integrates relative abundances of taxa with the ecological distances between habitat clusters they were observed in. At the species level, the generalism score ranges from 0 for species predominantly abundant in a single habitat cluster (indicative of specialization^49^) to 0.512 for *Afipia sp000497575*, a species thriving across disparate habitat clusters at comparable relative abundances (**Figure 4A, B, Methods, Supplementary Figure 9A, B, Supplementary Table 3**; Generalism scores for all taxa are accessible on the SPIRE website (https://spire.embl.de/)).

**Figure 4.**
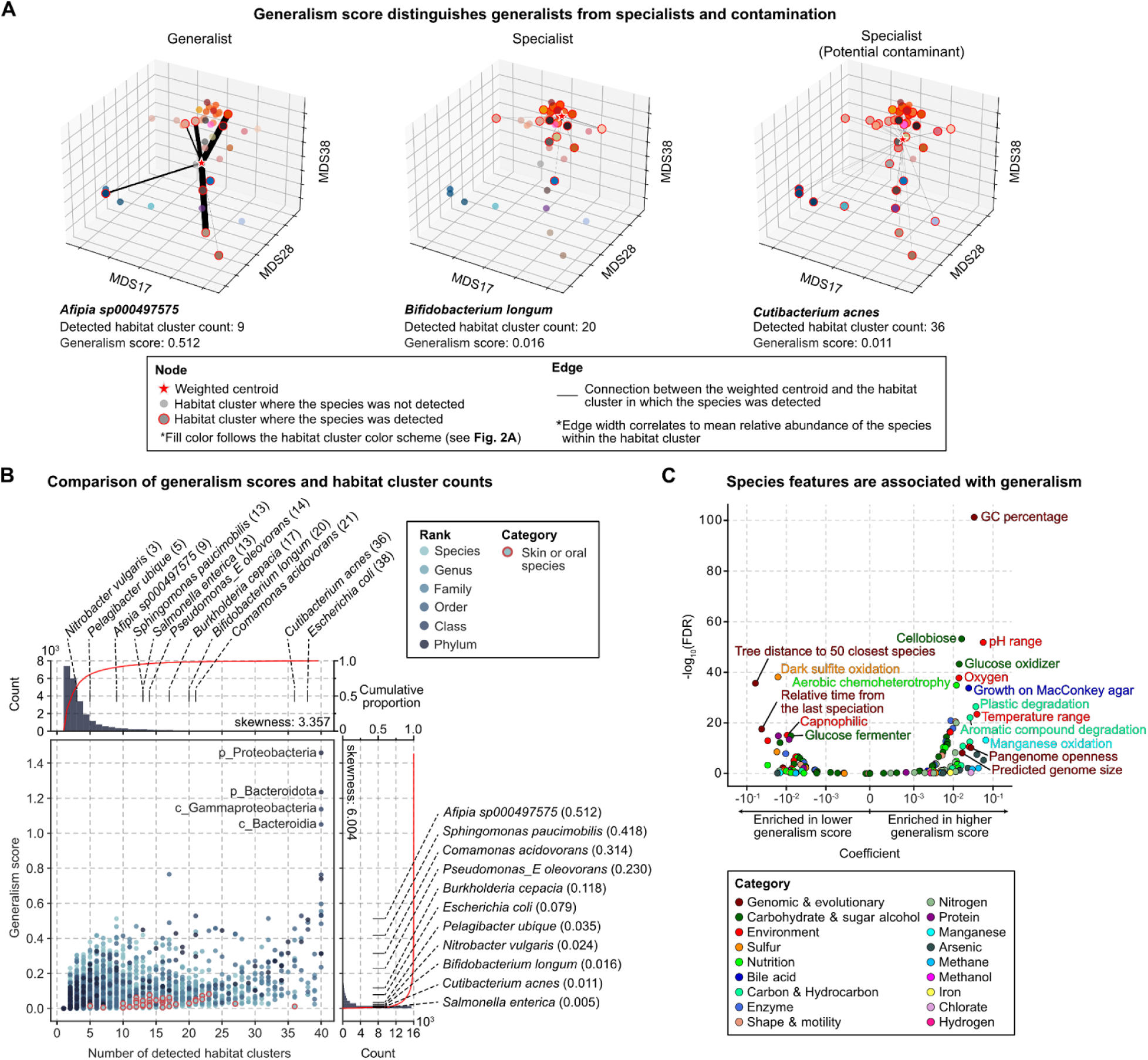
Prokaryotic generalism score and traits enriched in generalists. **(A)** Illustration of the integrative generalism scores for *Bifidobacterium longum*, *Afipia sp000497575*, and *Cutibacterium acnes*. *B. longum* was detected in 20 animal gut clusters (including human), yet these clusters were ecologically similar, resulting in a low generalism score (0.016). In contrast, *Afipia sp000497575* was detected in fewer habitat clusters (9 clusters) and showed consistent abundance across ecologically distinct habitats (including aquatic, terrestrial, and bovine gut clusters), yielding a higher generalism score of 0.512. A potential contaminant species, *C. acnes*, was detected in 36 habitat clusters but had disproportionately high abundance in human skin-associated (HSS) clusters, resulting in a low generalism score of 0.011. **(B)** Comparison of generalism scores and habitat cluster counts. Histograms and cumulative proportions (red lines) are shown on the upper panel (for cluster count) and the right panel (for generalism scores). Species frequently (> 50%) found in skin and oral samples are highlighted in red. Raw habitat cluster counts can inflate generalism predictions when taxa are frequent sample contaminants (e.g., skin or oral species). Skewness was calculated using Fisher’s moment coefficient. **(C)** Volcano plot representing species traits associated with prokaryotic generalism. Positive coefficients indicate traits enriched in generalist species, while negative coefficients represent traits enriched in specialist species. GC content is the trait most positively associated with prokaryotic generalism. FDR was calculated by adjusting the *p*-values from the logistic regression analysis using the Benjamini-Hochberg multiple testing correction.

Compared to simply counting the number of habitat clusters in which a species was detected^50^, our scoring method yielded lower estimates of generalism (Fisher’s moment coefficient = 6.004 versus 3.357). For simplicity, we operationally define species with a generalism score below 0.1 as specialists, which includes 95.4% of all species, and those with a score of 0.1 or above as generalists (**Figure 4B**). This aligns with previous reports that most microbes are specialists, while only a small fraction are generalists capable of colonizing highly disparate habitats^4,5^. Identifying and quantifying generalism can be biased by factors such as human-derived contamination, where oral and skin microbes are detected in environmental samples and may be misinterpreted as evidence of broad adaptability (for which we find indications, see **Supplementary Figure 6B, C**). However, by taking the species relative abundance into account, our scoring scheme appeared robust against such biases (**Figure 4B**). For instance, *Cutibacterium acnes*, a known skin commensal and a contamination indicator of metagenomic samples^51^, was detected in 36 out of 40 habitat clusters. Due to its abundance pattern with disproportionate preference for the HSS cluster, *Cutibacterium acnes* was not misinterpreted as a generalist (generalism score = 0.011, **Figure 4A, B**).

We identified traits, pathways, and genomic features associated with generalism using linear regression analysis, controlling for genome completeness and contamination. To focus on features consistently enriched among generalists across diverse prokaryotic lineages, we also accounted for phylogenetic effects (**Methods**). The generalism score showed strong positive associations with a broader tolerance range for both temperature and pH, which suggests environmental flexibility (linear regression FDR-corrected *p (q)* < 1e-20) (**Figure 4C**). These associations were more pronounced than those based on simple habitat cluster counts (**Supplementary Figure 10A**). We also observed a marked positive correlation between generalism and GC content (linear regression *q* < 1e-100) (**Figure 4C, Supplementary Figure 10B**), consistent with previous findings linking higher GC content to resource opportunism, a key factor in generalism^52^. This correlation is reflected in the high fraction of generalists in terrestrial and anthropogenic habitat clusters, whose microbiomes exhibited high GC contents (**Figure 3C**). Notably, generalism was associated with higher speciation rates (**Figure 4C**, specialists showed larger phylogenetic distances to their closest species and longer intervals since their last speciation, linear regression *q* < 1e-15), consistent with previous reports^50,53^.

Traits associated with oxygen utilization (e.g., aerobic chemoheterotrophy, glucose oxidation, and aerobic respiration) were positively correlated with generalism (linear regression *q* < 1e-20). In contrast, traits linked to oxygen-deficient or high-CO_2_ environments (e.g., capnophilic metabolism, glucose fermentation, and dark sulfite oxidation) were negatively correlated with the generalism score (linear regression *q* < 1e-10) (**Figure 4C**). This suggests that aerobic respiration supports higher generalism, aligning with the oxygenated nature of most habitats. Furthermore, traits and pathways involved in degrading diverse chemical compounds were enriched in generalists (**Figure 4C**, **Supplementary Figure 10C**), highlighting the importance of resource opportunism. Specialists, on the other hand, were enriched for pathways associated with antibiotics and polyketide biosynthesis, secondary metabolites commonly used for antibiotic production (**Figure 4F**, **Supplementary Figure 10C**). This implies that antibiotic production may increase fitness for specialists by competing with sympatric species^54^, consistent with the competition-dispersal trade-off concept in ecology^55^.

### Generalists mediate horizontal gene transfer between ecologically disparate habitats

Due to their adaptability to diverse environmental conditions and resources, generalist microbes thrive across a broad range of habitats. Through dispersal, generalists can transport both their own genetic material and horizontally acquired genes (HGT) from sympatric species into new habitats (**Figure 5A**). Because species co-occurrence increases the likelihood of horizontal gene transfer^56^, and generalists can be members of microbiomes from different habitats, generalists have the potential to be key facilitators of planetary gene flow, particularly across ecologically disparate habitats. To quantify and validate this hypothesis, we screened ∼2 million genomes and MAGs for high-confidence HGT events, adapting methods from Khedkar et al., 2022^57^ (species sharing five adjacent genes, including a recombinase, **Methods**). We integrated these HGT events with our habitat cluster and generalism framework, normalizing for sampling biases among habitat clusters (**Methods**).

**Figure 5.**
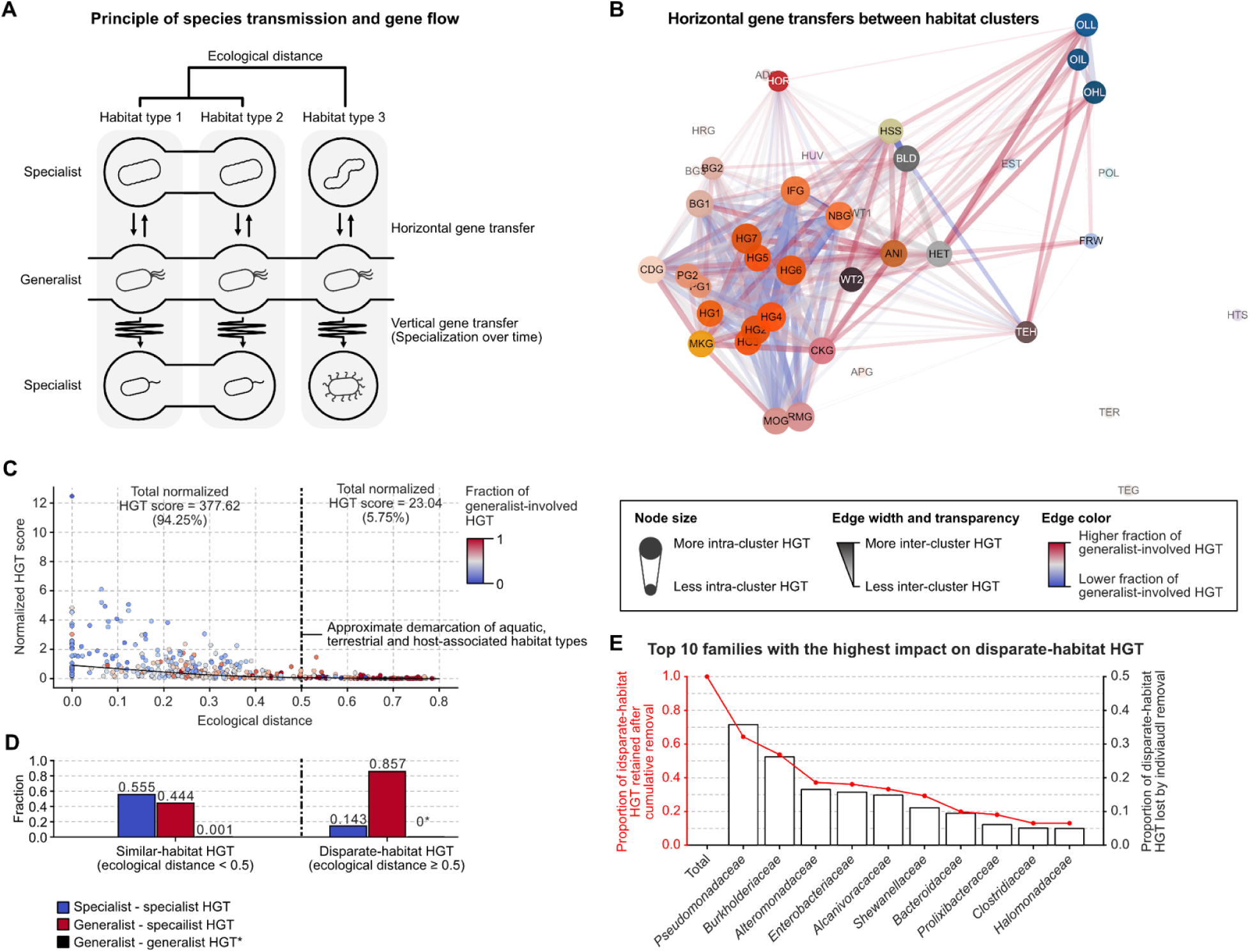
Generalists as a vehicle for HGT across ecologically disparate habitats. **(A)** A conceptual visualization of possible gene flow between habitats, by microbial dissemination or horizontal gene flow between microbes. Generalists, but not specialists, can transport their own or horizontally acquired genes into new disparate habitats. **(B–D)** Generalists are involved in the majority of HGT events between disparate habitat clusters. **(B)** Network visualization of horizontal gene transfer between habitat clusters. The position of the nodes (habitat clusters) was determined based on shared taxa (Figure 2A). Edges represent horizontal gene transfer events between microbes detected in different habitat clusters. The edge width and transparency scale with the count of HGT events between habitat clusters (normalized by the count of assembled MAGs per habitat cluster). Edge colors indicate the fraction of generalists that are involved in the HGT events. Habitat clusters with < 1,000 MAGs with > 50% completeness were not considered for the HGT analysis (transparent nodes). **(C)** Scatter plot visualization of normalized HGT score along ecological distances. The point color scales with the fraction of generalists that are involved in the HGT events, in analogy to the edge colors in (B). The vertical dashed line marks an ecological distance of 0.5 as operational definition of disparate habitat clusters. Smooth line represents a lowess local regression. **(D)** Proportion of HGT events between specialists and generalists within short ecological distances (< 0.5) or across disparate habitat clusters (> 0.5 ecological distance). **(E)** Families involved in most disparate-habitat HGT events. The top three families are jointly involved in 63% of HGT events across disparate habitat clusters.

Most of the 84,049 HGT events detected this way occurred between ecologically similar habitat clusters, such as within host-associated or ocean clusters (**Figure 5B, C**). Although specialists were involved in over 99% of all HGT events, the fraction of generalist-involved HGT increased significantly with greater ecological distance between the sampled environments (Pearson’s *r* = 0.589, *p* = 1.82e-35, **Supplementary Figure 11**). Specifically, at an ecological distance threshold of 0.5 as an operational separation of similar and disparate habitat clusters (differentiating aquatic, terrestrial and host-associated habitats, **Figure 2A**), similar-habitat HGT accounted for 94.25% (**Figure 5C**) of all HGT events, of which 55% occurred exclusively between specialists. In contrast, among disparate-habitat HGTs (5.75% of events), only 14.3% involved specialists exclusively, whereas 85.7% involved interactions between both specialists and generalists (**Figure 5D**). Thus, despite being rare, HGT between disparate habitat clusters was disproportionately mediated by generalist taxa. Notably, species from only three prokaryotic families, *Pseudomonadaceae*, *Burkholderiaceae* and *Alteromonadaceae*, together contribute to the majority (63%) of disparate-habitat HGT (**Figure 5E**). This suggests that targeted measures aimed at interrupting gene flow between disparate gene reservoirs (e.g., antibiotic resistance genes) could effectively focus on these key taxa.

These findings uncover fundamental principles of planetary-scale gene flow, highlighting generalists as key vehicles capable of overcoming ecological and geographical boundaries, thereby facilitating HGT across ecologically distinct microbial habitats.

### Generalists shape anthropogenic microbiomes and carry antibiotic resistance genes to disparate habitats

Having identified generalist microbes as major vehicles of HGT across ecologically disparate habitats, we next explored their role in disseminating antibiotic resistance genes. Notably, disparate-habitat HGT events often occurred via anthropogenic interfaces, such as wastewater (WT2), built environments (BLD), and surfaces frequently exposed to human contact (HET, **Figure 5B**). Within the planetary microbiome landscape, these habitats exhibit a unique pattern: taxonomically, they resemble host-associated microbiomes, yet functionally they share traits with environmental microbiomes (**Figure 3** - dashed rectangle ii and iii, **Figure 6A**). Importantly, they also harbor a high relative abundance of generalist species (**Figure 3C**; BLD: 7.74%, WT1: 7.92%, WT2: 11.33%, HET: 10.65%; higher than all habitat clusters but TER: 13.86%, TEH: 13.61%), suggesting an advantage for generalists in these environments. We therefore hypothesized that adaptive generalist taxa from human microbiomes disseminate and integrate into anthropogenic microbiomes, structurally shaping the local microbial communities and facilitating genetic connectivity between disparate habitats.

**Figure 6.**
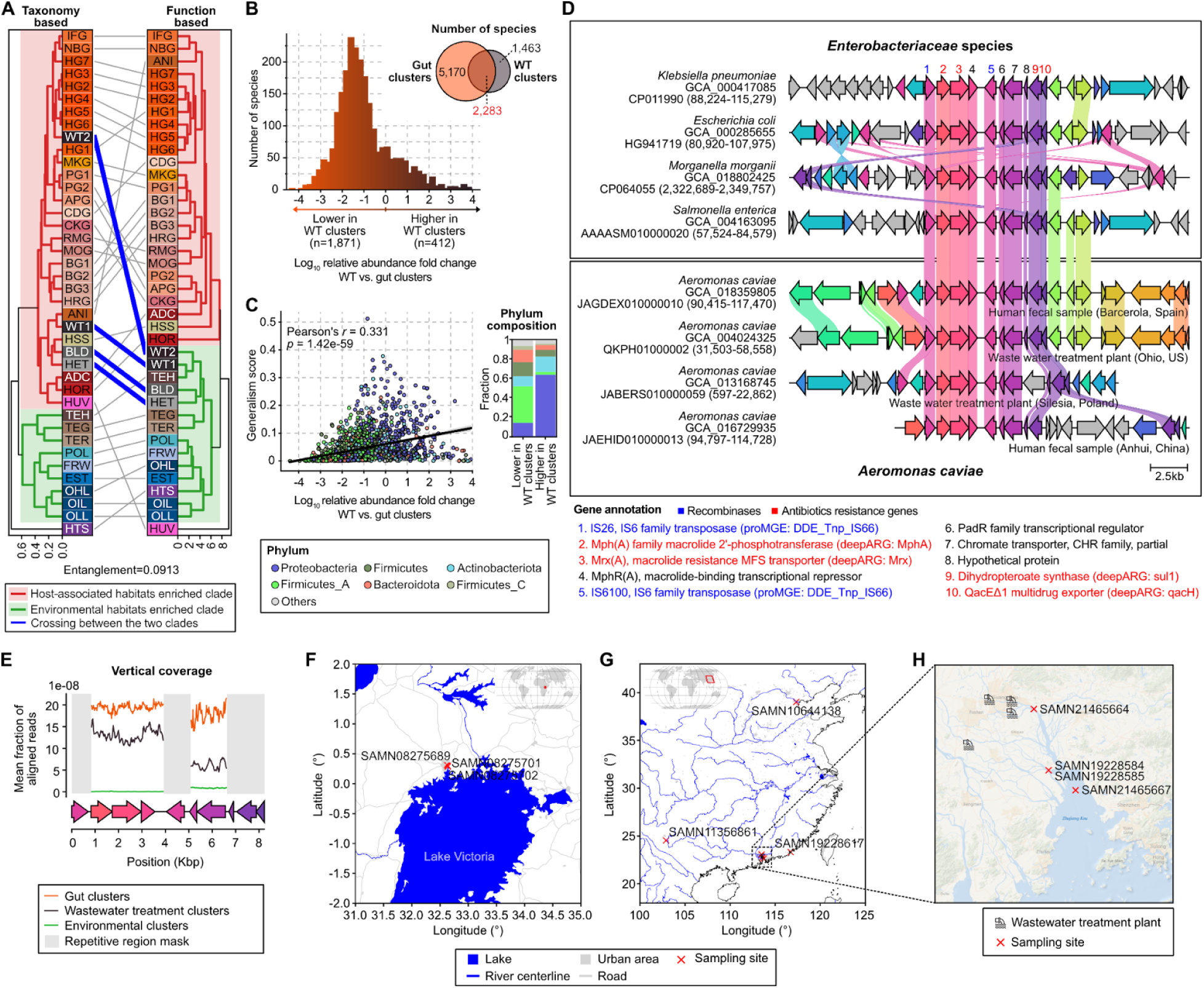
Generalists transport antibiotic resistance genes across ecological and geographical boundaries. **(A)** Tanglegram comparing the placement of habitat clusters in the taxonomic (left) versus functional (right) microbiome landscape (hierarchical clustering). Grey lines connect habitat clusters which were placed in the same branch when comparing the taxonomic and functional microbiome landscape. Bold blue lines highlight habitat clusters which are taxonomically placed in the host-associated branch (red) but functionally in the environmental branch (green), which exclusively marks anthropogenic habitats. **(B)** Venn diagram showing the number of species detected in gut and wastewater clusters (top-right). The histogram represents the distribution of the relative abundance fold change for 1,864 overlapping gut/wastewater species. Positive abundance change indicates higher relative abundance in wastewater, compared to gut clusters. The majority of taxa detected in wastewater clusters were also found in gut microbiomes (60.95%) but frequently (81.95% of cases) at a lower relative abundance. **(C)** Generalism scores are positively correlated with the relative abundance change of taxa in wastewater (compared to gut clusters). Data points are colored by the phylum of each species. Barplot represents phylum composition of species with increased or decreased relative abundance in wastewater, compared to gut clusters. Proteobacteria species exhibited increased relative abundance in WT clusters compared to gut clusters, whereas Firmicutes_A species showed a decrease. **(D)** Evidence of a potential antibiotic resistance island consisting of 10 genes, shared between gut specialist *Enterobacteriaceae* species (showing 4 out of 36 genomes) and the generalist *Aeromonas caviae* (showing 4 out of 13 genomes). Genes with 100% sequence identity are shown in identical colors. The annotations for these genes are provided below, with blue text indicating insertion sequences, and red text indicating antibiotic resistance genes. **(E)** Vertical coverage of the potential antibiotic resistance island region (panel D) by metagenomic reads summarized by habitats. The genomic regions that are found repeatedly across prokaryotic genomes (**Supplementary Figure 12E**) are masked. The potential resistance island was mostly detected in gut and wastewater habitat clusters and rarely in environmental samples. **(F–G)** Geographical locations (marked by red crosses) of environmental samples at (F) Lake Victoria near Kampala and (G) sampling points in China where the resistance island was detected. **(H)** Map highlighting the Guangdong-Hong Kong-Macao Greater Bay Area, along with the sampling points and locations of nearby wastewater treatment plants in the region.

To test this, we specifically focused on wastewater clusters (WT1–2), given their high and continuous influx of host-associated microbiota and their well-established role as hubs for the selection^58,59^ and horizontal transfer of antibiotic resistance genes^60^. A substantial fraction of species (60.95%, 2,283 out of 3,746) identified in the WT clusters were also present in animal (including human) gut clusters (HG1–7, NBG, IFG, MOG, RMG, CKG, APG, MKG, PG1–2, CDG, BG1–3, HRG), in line with previous research^61,62^. The vast majority of these shared species (81.95%, 1,871 out of 2,283) exhibited lower relative abundance in wastewater compared to their respective gut clusters (**Figure 6B**), consistent with an enrichment of traits characteristic of anaerobic specialization (e.g., anaerobic metabolism, glucose fermentation; logistic regression *q* < 1e-30, **Supplementary Figure 12**). Species exhibiting higher relative abundance in wastewater, in contrast, demonstrated a significant enrichment of traits associated with adaptation to aerobic conditions (e.g., aerobe, oxidase activity, glucose oxidation; logistic regression *q* < 1e-30; Methods, Supplementary Figure 12). Importantly, we found a significant positive correlation between relative abundance fold-change from gut to wastewater habitats and generalism scores (Pearson’s *r* = 0.331, *p* = 1.42e-59, **Figure 6C**), supporting our hypothesis of a selective enrichment of human-derived generalists in wastewater, where they contribute to shaping the local microbial communities.

Microbiological studies of wastewater have been extensively conducted, as effluents, despite processing, can still carry microbes, including pathogens, into freshwater systems^63,64^. To study to what extent generalists facilitate the dissemination of genes beyond wastewater, we closely examined the ten species that are most strongly associated with disparate-habitat HGT events, specifically focusing on the transfer event with the most antibiotic resistance genes to further illustrate broader ecological and public health implications (i.e., multi-drug resistance). This revealed an antibiotic resistance island with ten genes, including four putative ARGs, that was shared between the generalist *Aeromonas caviae* and multiple *Enterobacteriaceae* gut specialists (**Figure 6D, Supplementary Figure 13B, Methods**). This island also harbored the highest amount of ARGs horizontally transferred between host-associated and anthropogenic environments in a single event. *A. caviae* are recognized for their robustness to wastewater treatment procedures^65^, and are classified by the US Environmental Protection Agency as opportunistic pathogens and potential water contaminants^66,67^. Within shared coding regions, the identified resistance island had complete sequence identity across all carrier species, indicating recent horizontal transfer, which is in alignment with the emergence of wastewater treatment plants as hubs of such gene exchange^60,68^.

The resistance island was most abundant in gut and wastewater samples (**Figure 6E, Supplementary Figure 13C, D**), but notably also appeared in estuary samples downstream from wastewater facilities (**Supplementary Figure 13C**). We mapped reads from *A. caviae*-positive metagenomes to the resistance island (**Methods**) and detected it in 10 environmental samples. These environmental samples were likely impacted by anthropogenic activities: Lake Victoria near Kampala, where untreated wastewater was discharged into the lake^69^; biofilms on plants irrigated by Haihe River water; Fuxian Lake near Kunming; and estuarine sites downstream of wastewater treatment plants (**Figure 6F–H**), within the densely populated metropolitan region Guangdong-Hong Kong-Macao Greater Bay Area. Although directionality and causality still need to be shown, our results suggest that generalists such as *A. caviae* are capable of facilitating the transfer of ARGs between host-associated and environmental microbiomes. Our habitat and generalism framework thus provides a powerful approach for tracing HGT (including antibiotic resistance spread) across habitats at a planetary scale.

## Discussion

In this study, we leveraged a planetary-scale metagenomic dataset^24^ to develop and apply a robust and fine-grained microbiome-based habitat clustering framework. Our framework harmonizes dispersed knowledge from various ecosystem-specific studies, allowing us to quantify the ecological drivers, functional traits, and adaptation strategies of microbiomes that emerge across their ecological distances. By developing a scoring scheme to characterize the continuum from prokaryotic specialists to generalists, quantifying their dispersal between habitat clusters, and integrating horizontal gene transfer events detected in our dataset, we identified a fundamental principle that underlies the growing importance of generalists as mediators of gene flow with increasing ecological habitat distance.

Our study establishes a model that captures the planetary-scale microbial metacommunity^70^, thereby extending ecological frameworks, previously well-established for animals^71^ and plants^72,73^ to prokaryotes. The metacommunity view is a prerequisite for understanding how the planetary microbiome’s structure and function are shaped and how they evolve: Our findings reinforce the classical microbiological tenet formulated by Baas Becking that “everything is everywhere, but the environment selects”^74,75^. This is evident through the clustering of metagenomes by habitat conditions rather than geography, best illustrated by the similarity of ocean microbiomes towards polar regions (**Figure 2E**) and by previous reports on terrestrial and lacustrine microbiomes from the Arctic and Antarctica^39^. We note that the Baas Becking tenet primarily applies to specialists, whereas generalist species deviate because their pronounced adaptability allows them to transcend ecological constraints. Furthermore, we propose specific mechanisms underlying gene flow across disparate habitats, notably mediated by HGT events between specialists and generalist species, as exemplified by the spread of antibiotic resistance genes between gut, wastewater, and other aquatic habitats.

While our study synthesizes various concepts from molecular ecology and evolution and is supported by established knowledge, our approach has limitations. For example, sampling imbalances in public research are also reflected in our dataset, impacting both the accuracy and sensitivity of microbial profiling tools across habitats and limiting the availability and resolution of taxonomic profiles for cluster formation. Moreover, our analysis focused exclusively on prokaryotes, omitting eukaryotic microorganisms and viruses, despite their well-documented importance in shaping microbial communities^76,77^. Including these organisms, although technically challenging, could redefine cluster boundaries and uncover finer-scale ecological patterns. Lastly, as a cross-sectional study, our research does not capture evolutionary processes such as speciation or long-term adaptation, which would require paleogenomic data spanning evolutionary timescales^78^ and is beyond the scope of this work.

Future research could build upon our planetary habitat clustering framework to monitor and quantify microbial gene flow in response to changes in environmental and anthropogenic pressures. As datasets continue to grow, this foundational work paves the way to predict microbiome structure from habitat descriptors and to better understand and manage human impacts on the global dissemination of microbial genes with relevance for planetary health.

## Methods

### Taxonomic, functional and k-mer-based distance between metagenomic samples

We compiled an mOTU v3.0.1^79^-based taxonomic profile for 85,604 metagenomic samples collected from 690 studies worldwide using SPIRE^24^. Samples with < 100 mapped reads or more than 70% of their profile unclassified (indicated as ’-1’) were excluded to maintain data integrity. We removed the unclassified feature from the profiles and renormalized each profile so that the sum of sample values equals one. The mOTUs profiles were converted to the GTDB r207 taxonomy^80,81^, and we generated profiles at different phylogenetic ranks, from family to species. Additionally, we created concatenated profiles spanning from domain to species. We applied arcsine and log transformations to the relative abundance profiles. The arcsine transformation involved taking the square root of the profile values and then calculating the arcsine. For the log transformation, we added half the global minimum value to the profile, took the natural logarithm, and adjusted the transformed values to range between 0 and 1.

For each sample, functional profiles on different hierarchies were gathered. From assemblies with a length of > 1,000 bp, open reading frames (ORFs) were predicted using Prodigal^82^ (v2.6.3) and functionally annotated with eggNOG-mapper^83^ (v2.1.3). To approximate ORF abundances, each ORF was multiplied by the average sequencing coverage of the contig it was found on. Then, the counts were summed on the single-letter Cluster of Orthologous Groups (COG) category level and on the root-level COG annotation. A sample-wise compositional transformation was applied, yielding relative abundance values per functional category. To generate a functional profile on the level of single-letter COG categories per habitat cluster, proportions from all samples from the same habitat cluster were summed and transformed to cluster-wise relative abundances.

For the taxonomic (mOTUs level) and functional (root-level COG) profiles, we calculated pairwise Bray-Curtis dissimilarity for all sample pairs. Additionally, we combined the functional profile-based distance matrix with the taxonomic profile-based distance matrices derived from mOTUs, GTDB-species, and GTDB-domain-to-species. The combined distance matrices were generated using varying ratios, ranging from 1:9 to 9:1, in increments of 1. We also calculated 21-mer overlap based distance between raw sequencing reads of the metagenomic samples using Mash^30^ (v2.3).

### Dimensionality reduction and clustering of metagenomic samples

For each distance matrix, we applied dimensionality reduction techniques using UMAP^28^, DensMAP^84^, and t-SNE^85^. For UMAP and DensMAP, we experimented with both 3-dimensional and 2-dimensional configurations, varying the *‘n_neighbors’* parameters (16 levels: 10, 15, 20, 25, 30, 35, 40, 45, 50, 55, 60, 65, 70, 80, 90, 100) and *‘min_dist’* parameters (4 levels: 0, 0.01, 0.05, 0.1). For t-SNE, we also used both 3-dimensional and 2-dimensional configurations, adjusting *‘early_exaggeration’* (4 levels: 12, 24, 36, 48, 100) and *’perplexity’* parameters (9 levels: 5, 10, 15, 20, 30, 40, 50, 75, 100).

Utilizing the coordinates obtained from each dimensionality reduction space, we then clustered the samples using HDBSCAN^86^, k-means, and hierarchical clustering methods. For HDBSCAN, we tested various *‘min_cluster_size’* (5 levels: 10, 50, 100, 150, 200) and *‘min_samples’* (7 levels: 1, 5, 10, 20, 50, 75, 100) parameters. Hierarchical clustering was performed using Euclidean distance-based average linkage. For both k-means and hierarchical clustering, we tried multiple *‘n_clusters’* (41 levels: every 5 from 10 to 200, 250, and 300) parameters. We observed that t-SNE underperforms compared to UMAP and densMAP for our habitat clustering purpose. Therefore, we skipped t-SNE for the function-taxonomy combined distance matrix to reduce the computational burden.

As a result, a total of 1,658,865 clustering models were created by combining different profile pre-processing methods, dimensionality reduction algorithms, and clustering methods with many of their parameters (**Supplementary Table 1**).

### Benchmarking of clustering results and selection of habitat clusters

For each clustering model, we calculated the Shannon entropy for each cluster based on a simplified microntology (Figure 1). Clusters composed exclusively of samples originating from a single microntology had an entropy value of zero, whereas clusters comprising samples from heterogeneous microntology terms exhibited higher entropy values. We used the mean entropy score as the representative entropy value for each clustering model. Additionally, we calculated the silhouette index based on Euclidean distances in the UMAP embedding for each clustering model. We conducted a comprehensive comparison across various features (taxonomy, function, combined taxonomy-function, k-mer), transformation methods (raw, log, arcsine), dimensionality reduction methods (UMAP, densMAP, t-SNE), and clustering methods (HDBSCAN, hierarchical, K-means), exploring a range of dimensionality reduction and clustering parameters. From this analysis, we observed that clustering based on 3D UMAP resulted in lower entropy and higher silhouette index values compared to other dimensionality reduction methods. Therefore, among the clustering models based on 3D UMAP, we selected the model (log-transformed species-level GTDB taxonomic profiles, Bray-Curtis dissimilarity, 3D-UMAP with n_neighbors=25 and min_dist=0.01, hierarchical clustering with n_clusters=105, Supplementary Figure 4) with the lowest mean entropy that also satisfied the following criteria: silhouette index in the top 10% (> 0.511), mean entropy in the bottom 10% (< 0.306), < 100 clusters, and > 95% of samples included in clusters with a size of ≥ 100.

If a cluster consists primarily of samples from a single study, it is difficult to determine whether it represents a unique study protocol or the environmental characteristics of the habitat where the samples were collected. Therefore, to select clusters that genuinely represent a habitat type rather than study-specific bias, we analyzed the associations among the number of samples, the number of unique studies from which the samples were derived, and the fraction of samples derived from the most dominant study. As expected, smaller cluster size, fewer unique studies, and higher dominance by a single study were correlated. Therefore, we excluded clusters that had 100 or fewer samples, clusters derived from two or fewer studies, and clusters in which more than 95% of samples were derived from a single study. This process retained 39 out of the 105 clusters (**Supplementary Figure 3A**–C).

Among the 39 clusters obtained, we manually divided a cluster, APG-HRG, into two subclusters, APG and HRG. This decision was prompted by the observation that cluster 13 was clearly separated into two distinct groups in the UMAP embedding (**Supplementary Figure 3D**), with any pairwise distance between samples from different subclusters being greater than any pairwise distance within the same subcluster (**Supplementary Figure 3E**). Additionally, we confirmed that the distribution of Bray-Curtis distances significantly decreased after dividing into subclusters, indicating increased homogeneity within each group (**Supplementary Figure 3F**). Furthermore, both subclusters comprised animal gut microbiome samples; however, subcluster APG was predominantly composed of samples from apes (*Gorilla* and *Pan* species), while subcluster HRG was mainly composed of horse samples (**Supplementary Figure 3G, H**). Although these subclusters were not defined by the initial systematically benchmarked clustering parameters and their sizes were < 100 samples, we concluded that dividing and analyzing this cluster based on their clearly distinguishable characteristics would better define clusters that reflect microbiome habitats. Therefore, we used these subclusters as the final habitat clusters. Through this process, we ultimately defined 40 habitat clusters consisting of a total of 71,871 samples (**Supplementary Table 2**).

### Ecological distance between habitat clusters

The dimensionality reduction algorithms employed in this study (UMAP, densMAP, and tSNE) prioritize the preservation of local structure over global structure. Therefore, the distances between clusters were calculated based on the distance from the original taxonomic profile, not from dimensionality reduction space.

For each habitat cluster 𝐶, we defined the set of detected taxa 𝑇 as the taxa whose prevalence exceeds a specified cutoff value. The overlap coefficient distance 𝐷 between two clusters 𝑐*_i_* and 𝑐*_j_* is defined as:

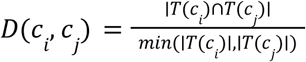

where 𝑇(𝑐*_i_*) and 𝑇(𝑐*_j_*) represent the sets of detected taxa for clusters 𝑐*_i_* and 𝑐_j_, respectively.

Different prevalence cutoffs have distinct implications. A higher prevalence cutoff emphasizes the overlap of core taxa between habitat clusters, while a lower prevalence cutoff accounts for the overlap of taxa that are sporadically present in the clusters. Rather than using a single prevalence cutoff, we calculated *D* across a range from 0.01 to 0.3, increasing by 0.01 at each step, and calculated the average. We evaluated the robustness of clades in the hierarchical dendrogram generated from these distances by assessing how consistently each clade is recovered across the different cutoffs. The average distance was computed at various taxonomic ranks: from species to phylum. Since differences at higher taxonomic ranks indicate more substantial divergence, we assigned weights to the rank-specific distances inversely proportional to the median GTDB Relative Evolutionary Divergence (RED) score^87^ at each rank (phylum:class:order:family:genus:species = 3.03:2.22:1.61:1.32:1.09:1). The final ecological distances between the habitat clusters were determined as the weighted average of these rank-specific distances.

### Network visualization

The network was visualized using Cytoscape v3.10.2^88^. In the network shown in **Figure 2A**, the Prefuse Force-Directed Layout^89^ was used. The weight for layout computation was set as the negative logarithm of the ecological distance. The layout parameters were set as follows: number of iterations = 50,000, default spring coefficient = 1, default spring length = 50, and default node mass = 5. The width and transparency of the edges were determined by the ecological distance between the connected habitat clusters (shorter distances resulted in thicker and more opaque edges). The node size was determined by the logarithm of the number of samples belonging to each habitat cluster.

In the network shown in **Figure 5B**, the layout determined in **Figure 2A** was preserved (maintaining the relative positions of habitat clusters based on ecological distance), and horizontal gene transfer (HGT) information was overlaid. In this network, node size represents the normalized score of intra-cluster HGT, while the edge width and transparency represent the normalized score of inter-cluster HGT. The edge color was determined by the fraction of generalist-involved HGTs among all HGT events between the two clusters. A total of 11 clusters were excluded (transparent nodes) from the network visualization because the small number of samples and resulting low number of assembled MAGs could introduce distortions in the calculation of normalized HGT scores.

### Analysis of host & human DNA contents

To detect human or host DNA contamination in metagenomic samples, we utilized the Sequence Taxonomic Analysis Tool (STAT)^90^ provided by the National Center for Biotechnology Information (NCBI) Sequence Read Archive (SRA)^91^. We focused on samples from the FijiCOMP^92^ project that lacked host annotations; samples annotated as ocean or freshwater in microntology and metalog but classified within the HSS cluster; and randomly selected samples from OIL, OLL, OHL, and FRW. For these samples, we compiled all associated run accessions and used the BigQuery UI to examine the taxa to which the reads were mapped. To identify the missing hosts in the FijiCOMP project samples, we investigated the most dominant vertebrate genus among those with more than 1,000 mapped reads. For assessing human DNA contamination, we calculated the proportion of reads mapped to the genus *Homo*.

### Cluster level summary of species’ genomic and phenotypic features

From the GTDB metadata, we selected up to 50 genomes with the highest completeness for each species. These genomes were chosen among isolated genomes and metagenome-assembled genomes (MAGs) with a CheckM2^93^ completeness > 50% and contamination < 10%. The estimated genome size for each genome was calculated as genome size × (100 / completeness). For each species, we used the mean GC content, mean coding density, and mean estimated genome size as representative values.

For pangenome openness, we compiled metagenome-assembled genomes (MAGs) and isolate-derived genomes with > 80% completeness and < 5% contamination, as assessed by CheckM2^93^, from the SPIRE^24^ and Progenomes3^94^ databases. For each GTDB species, we selected up to 500 genomes evenly from both databases. Species with < 10 genomes were excluded from pangenome openness calculations. Genes were predicted using Prodigal and clustered at 95% nucleotide identity using MMseqs2 linclust^95,96^ (--cov-mode 0, -c 0.95). Genomes were added incrementally, and unique gene clusters were counted. This process was repeated 50 times with randomized genome order, and the resulting rarefaction curve was fit to a power law equation (*y* = *kx*^α^). The power law exponent, α, from this model was used to quantify pangenome openness. A higher α indicates a more open pangenome, where new genes are continually discovered as more genomes are sampled^97^. The power law equation fits the data well across most species, with a median R-squared value > 0.997.

For evolutionary features, we utilized the phylogenetic tree provided by GTDB (release 207) to calculate the tree distance to the 50 closest species for each species. Additionally, we computed the Relative Evolutionary Divergence (RED) score for all species using PhyloRank^98^ (v0.1.12). For each species (leaf node), the RED score of its immediate parent node was used as the relative time since the last speciation event.

To quantify intra-species strain diversity, quality controlled metagenomic reads were aligned to species-specific markers (mpa_vJun23_CHOCOPhlAnSGB_202307) with MetaPhlAn ^99^. The resulting alignments were processed with SameStr ^100^ (v1.2024.02) to calculate the fraction of polymorphic sites for each species-level alignment, which served as a metric for intra-species population diversity as previously described^101^.

Phenotypic trait predictions were obtained for species in GTDB (r207) using MICROPHERRET^102^, Traitar^103^, GenomeSPOT^104^. We ran MICROPHERRET and Traitar MAGs and genomes from SPIRE^24^ and proGenomes3^94^. For this, we compiled 1,158,587 MAGs (> 50% CheckM2^93^-based completeness > 50%, contamination < 5%, and GUNC passed) from SPIRE and 907,388 genomes (completeness > 90%, contamination < 5%, GUNC passed) from Progenomes. We then binarized traits on a per-species level based on a 75% trait prevalence cutoff. GenomeSPOT predictions were taken directly from the manuscript^104^ and mapped from GTDB release r214 to r207 using GTDB’s metadata files.

We summarized all species features at the cluster level by calculating weighted averages using each species’ mean relative abundance within its habitat cluster as the weighting factor.

### Weighted centroid based generalism score

We computed the coordinates of each habitat cluster in a 40-component Multidimensional Scaling (MDS) space based on the ecological distances between habitat clusters. We confirmed that the Euclidean distances between habitat clusters in the MDS space exhibited a significantly high linear correlation with the phylogenetic distances (Pearson’s *r* > 0.97, *p* < 1e-300, **Supplementary Figure 9B**), indicating that the MDS representation accurately reflects the phylogenetic relationships among habitat clusters.

For each taxon 𝑡, we defined the set of habitat clusters, 𝐻, where the taxon is detected with the 1% prevalence cutoff. For each taxon 𝑡, we calculated the weight 𝑤*_t_*(ℎ), of the taxon for each habitat cluster ℎ, in which it was present. The weight was defined as the ratio of the taxon’s mean relative abundance, 𝐴, in the habitat cluster, to its mean abundance in the most abundant habitat cluster:

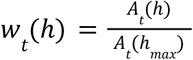

Where ℎ*_max_* is the habitat cluster where the taxon 𝑡 has the highest mean relative abundance. Therefore, the weight 𝑤_t_(ℎ) is proportional to the mean abundance of the taxon in the habitat cluster, ℎ, and is constrained to a maximum value of 1. We computed the coordinates of weighted centroid 𝐶*_t_* for each taxon in the MDS space as the weighted average of the coordinates of the habitat clusters where the taxon is present, weighted by 𝑤_𝑡_:

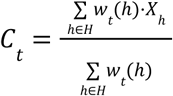

Where 𝑋*_h_* is the coordinate vector of habitat cluster ℎ in the MDS space. The generalism score 𝑆, for each taxon 𝑡 was calculated as the weighted sum of the squared Euclidean distances between the weighted centroid 𝐶*_t_* and the coordinates of the habitat clusters 𝑋*_h_* where the taxon is present:

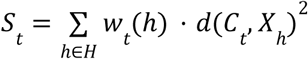

Where d represents the Euclidean distance function between the two input vectors. This score reflects both the distribution and abundance of the taxon across different habitat clusters. The scoring scheme is conceptually equivalent to weighted variance: the species’ weighted centroid corresponds to the mean, the clusters in which the species is present correspond to the observations, the Euclidean distance corresponds to the difference, and the relative abundance corresponds to the weight. According to our definition, taxa receive higher generalism scores if they are found in clusters that are more ecologically distant from each other, and if the taxon’s abundance is relatively uniform across the habitat clusters. (**Supplementary Figure 9A**).

### Species features associated with generalism score and abundance change in WT cluster

We summarized predicted traits (from Traitar^103^, MICROPHERRET^102^, and GenomeSPOT^104^), the presence and absence of KEGG Orthologous groups (predicted by EggNOG-mapper^83^), and other genomic features, such as genome length, pangenome openness, and tree distances to the 50 closest species, for each species (see **Methods**: **Cluster level summary of species’ genomic and phenotypic features** for details). All feature values were min-max normalized across species to standardize their ranges for linear regression. To prevent bias in the correlation analysis due to genome completeness and contamination, these factors were measured by CheckM2^93^ and included as covariates in the regression model. Since the range of generalism scores varies substantially across phyla, we accounted for phylogenetic effects by including binary covariates representing phylum membership in the linear regression model below:

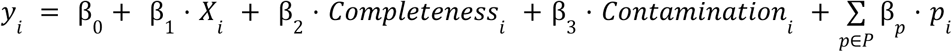

Where 𝑦_𝑖_ is generalism score for species 𝑖, 𝑋_𝑖_ is min-max normalized feature value for species 𝑖, and 𝑝_𝑖_ is binary value for phylum membership of species 𝑖. For the phylum-specific analysis, a similar linear regression model was applied, replacing phylum membership with class membership within the respective phylum.

To investigate the association between species traits and the abundance change in the WT cluster, we only used predicted traits from Traitar for simplicity. Logistic regression was performed to analyze the association between the predicted traits and abundance change in the WT cluster.

### Comparison between dendrograms

Taxonomy-based dendrograms were constructed by performing average linkage hierarchical clustering using the distance between clusters calculated based on the overlap coefficient as described above. We generated functional and trait-based dendrograms using average linkage hierarchical clustering based on the Euclidean distance between summarized root-level COG profiles (for functional) and resident species trait profiles (for trait-based) within each cluster. The entanglement between the different dendrograms was analyzed using the Python tenglegram package with an untangling process, which aims to find the pair of trees with minimized entanglement by flipping internal nodes while preserving the tree structures^105^.

### Identifying horizontal gene transfer with the context of generalism and ecological distance

Horizontal gene transfer (HGT) was detected using the method described by Khedkar et al. (2022)^57^ on SPIRE^24^ MAGs and Progenomes3^94^ isolated genomes. We used the same set of 20.7 million MAGs and genomes previously employed for phenotypic traits prediction. We clustered their genes with the mmseqs linclust^96^ using a 95% nucleotide identity cutoff with > 95% mutual alignment coverage (--search-type 3 --min-seq-id 0.95 --cov-mode 0 -c 0.95). If five or more consecutive genes appear in multiple species from different taxonomic families, and at least one of these genes encodes a recombinase, we define this as an HGT event. Because species from different families are phylogenetically distant, their vertically inherited genes (even highly conserved ones) typically do not exhibit more than 95% similarity. Thus, finding highly similar (> 95%) genes in species from different families indicates HGT. Moreover, if these highly similar genes occur consecutively in the genomes of both species and include recombinases, this further indicates that the genomic region has been horizontally transferred. We used a threshold of five consecutive genes because the median length of the smallest mobile genetic element type (IS_Tn, insertion sequences, and transposons) is approximately 5 kbp, which corresponds to roughly five prokaryotic genes.

To interpret HGT events in the context of prokaryotic generalism and ecological distances, we determined the ecological distance of each HGT event based on the habitat cluster information of the genomes or MAGs in which the event was detected. For MAGs derived from the SPIRE database, we used the habitat cluster of the sample from which the MAG was assembled. For genomes from the Progenome database, we identified MAGs with ≥ 99% genome ANI and used the habitat cluster information of the sample from which the matched MAG originated. If the HGT event occurred between a pair of habitat clusters with an ecological distance > 0.5, we classified this event as a disparate-habitat HGT.

Additionally, we classified each HGT event based on the generalism scores of the species involved. If all species involved in the event had a generalism score below 0.1, the event was classified as a specialist–specialist HGT. If at least one species had a generalism score of 0.1 or higher, the event was classified as a generalist-involved HGT. Among these, if all species had generalism scores ≥ 0.1, the event was further classified as a generalist–generalist HGT; otherwise, it was classified as a generalist–specialist HGT.

The number of detected horizontal gene transfer (HGT) events between habitat clusters increases with the number of assembled MAGs in each habitat cluster, as a higher MAG count provides more opportunities for HGT detection. Due to limitations caused by sampling bias, the number of samples and the number of assembled MAGs differ across habitat clusters. To address this bias, the HGT count between pairs of habitat clusters was normalized by dividing by the product of the number of MAGs assembled in the two clusters. Similarly, for intra-habitat cluster HGT, the count was normalized by the square of the number of assembled MAGs within the cluster. To minimize errors introduced by normalization, 11 habitat clusters (ADC, APG, BG3, EST, HRG, HTS, HUV, POL, TEG, TER, and WT1) with < 1,000 MAGs were excluded from the analysis.

### Identification of potential resistance islands and measuring the abundance from metagenomic samples

We annotated antibiotic resistance genes within the co-transfer using deepARG^106^. If a co-transfer contains antibiotic resistance gene(s), it is defined as a potential antibiotic resistance island. We focused on the horizontal gene transfer between *Aeromonas caviae*, a generalist species with a high proportion of disparate-habitat HGT and the most potential antibiotic resistance islands, and gut bacteria belonging to *Enterobacteriaceae*. This antibiotic resistance island was among the islands containing the greatest number of ARGs and was found in independently assembled, distinct genomes across multiple studies. The potential resistance island is visualized with the Clinker^107^.

To quantify the abundance of the potential resistance island across habitat clusters, we randomly selected 50 samples from each habitat cluster, for a total of 2,000 samples. Raw sequencing reads from these 2,000 samples were aligned to the resistance island region using Bowtie2^108^ (v2.5.2) with default parameters, and the number of aligned reads per DNA locus was determined using samtools^109^ (v 1.18) depth function. Vertical coverage was calculated as the number of aligned reads divided by the total number of reads, while horizontal coverage was calculated as the length of the aligned region divided by the total length of the resistance island. A horizontal-coverage threshold of 90% was applied to designate the resistance island as detected.

We observed that the regions from positions 0–850 bp, 3900–5100 bp, and 6600–8233 bp exhibited exceptionally high vertical coverage compared to the other regions (**Supplementary Figure 12D**). In particular, the 0–850 bp, 3900–5100 bp regions are aligned with insertion sequences, which are repetitive in the genome (**Figure 6D**). Therefore, we excluded these regions when comparing the relative abundance of the resistance island (**Figure 6E**) between different habitat clusters.

## Data and code availability

The underlying sequence data are available through the proGenomes (http://progenomes.embl.de/) and SPIRE websites (https://spire.embl.de/). Generalism scores for all taxa are accessible on the SPIRE website. The python scripts for the calculation of the generalism score and the horizontal gene transfer analysis are available at https://github.com/grp-bork/PlanetaryMicrobiomeHabitat_Kim_2025.

## Author contributions

Overall bioinformatics analysis: C.Y.K., D.P., and J.S., Conceptualization: C.Y.K., D.P., and A.O., Metadata curation: M.K., Data processing: A.F., M.K., C.S. and S.M.R., Horizontal gene transfer analysis: S.K., and A.G., Supervision: P.B., Writing manuscript: C.Y.K., D.P., J.S., and P.B.

## Declaration of generative AI and AI-assisted technologies in the writing process

During the preparation of this work the authors used ChatGPT (OpenAI) to improve language and clarity. After the usage of this tool, the authors reviewed and edited the contents as needed and take full responsibility for the content of the article.

## Acknowledgements

This work was supported by the National Research Foundation of Korea (NRF) grant funded by the Korean government (MSIT) (No. RS-2023-00240807), by the Deutsche Forschungsgemeinschaft (DFG, German Research Foundation) - project number 460129525 (NFDI4Microbiota), by the Ministry of Science, Research and the Arts Baden-Württemberg (MWK) within the framework of LIBIS/de.NBI, and by the Federal Ministry of Research, Technology and Space in the frame of de.NBI & ELIXIR-DE (W-de.NBI-014). The authors acknowledge the support of EMBL IT Services HPC resources (https://doi.org/10.5281/zenodo.12785829) for providing computational resources. The authors thank the Bork group and Thomas Sebastian Benedikt Schmidt for helpful discussions. We furthermore thank members of the von Mering group at the University of Zürich (João Frederico Matias Rodrigues, Janko Tackmann, Christian von Mering) for critical reading of the manuscript before submission.

