## Supplementary Information for "Planetary microbiome structure and generalist-driven gene flow across disparate habitats"

### Aquatic samples collected from transitional environments and extreme temperatures form distinct clusters.

Except for the groundwater samples, the majority of lentic and lotic freshwater samples clustered into the freshwater (FRW) cluster. However, samples collected from estuarine environments, such as the Savannah River Estuary in eastern USA, the Mississippi Sound in southern USA, and the Pearl River Estuary in southern China, formed a distinct cluster labeled EST (Estuary). Additionally, samples from hot springs in geographically diverse regions, including Yellowstone (USA), Hawaii (USA), Kootenay Rockies (Canada), Tshipise hot spring (South Africa), and Tengchong (China), clustered together in a separate designated Hot spring (HTS) cluster. Samples collected from Arctic and Antarctic lakes or ice sources also formed an independent cluster, Polar lake (POL). Although each of these clusters (EST, HTS, and POL) contains fewer than 200 metagenomic samples, the clustering of samples from geographically distant locations based on environmental similarities suggests that estuarine environments, as transitional zones, and extreme temperature environments such as hot springs and polar lakes support distinct microbial communities.

### The hydric status affects terrestrial microbiome composition

While limited metadata on soil hydric conditions makes it difficult to draw definitive conclusions, an examination of the environmental metadata for clustered soil samples suggests that soil samples may primarily be distinguished by the degree of hydric conditions. Most soil samples were classified into either the terrestrial hydric (TEH) or terrestrial (TER) clusters. The TEH cluster, in particular, included soil samples from boundary environments with aquatic characteristics, such as groundwater, aquifers, and sediment, which are classified as aquatic in microntology but are taxonomically close to terrestrial environments. Additionally, this cluster contained a large proportion of soil samples from wetlands, subsurface environments, and some sludge samples from wastewater treatment facilities. In contrast, the TER cluster was predominantly composed of samples from environments that could be considered less hydric, though not necessarily dry, such as forests, rhizospheres, arid regions, and tundra. The terrestrial grassland (TEG) cluster contained 180 out of 182 samples sourced from grassland environments. Although these samples were collected across three different studies (PRJNA405447, PRJNA365981, PRJNA449266) and over different years, they all originated from the same specific location: the Angelo Coast Range Reserve in California, USA. This unique clustering warrants careful interpretation, with the caveat that it is unclear whether it reflects general grassland environments, the specific soil characteristics of this region, or some other shared environmental variable.

### Differentiation of ancient and modern oral microbiomes

Samples collected from the human oral cavity, primarily saliva or dental plaque, were mostly classified into the Human oral (HOR) cluster. Interestingly, samples from multiple studies analyzing dental plaque of ancient humans formed a distinct ancient dental calculus (ADC) cluster. The microbial species found in the ADC cluster were a subset of those observed in the HOR cluster, with 96.6% of species prevalent more than 5% in the ADC cluster were also prevalent in HOR cluster (**Supplementary Figure 14A**). However, the prevalence and relative abundance of detected taxa was substantially different between the clusters (**Figure 3A**, **Supplementary Figure 14B**). A notable difference between ADC and

HOR was an archaeal species, *Methanobrevibacter_A oralis*, which was detected in 87% of samples in the ADC cluster but only in 0.6% of samples in the HOR cluster (**Supplementary Figure 14B**), consistent with previous studies[1,2]. As suggested by the studies, this reflects both shifts in oral microbiota composition over time between ancient and modern humans, as well as changes due to taphonomic processes that influenced microbial survival and abundance, contributing to the observed separation of the two clusters. Nevertheless, the HOR and ADC clusters remained the ecologically closest to each other, forming a robust clade (**Figure 2A**-clade e).

#### Human skin microbiomes are a major source of anthropogenic surface microbiomes

The Human skin and surface (HSS) cluster contains most human skin samples as well as a substantial number of metagenomic samples collected from built environment surfaces, such as hospital surfaces (PRJEB13117) and subways (PRJNA413474) (**Supplementary Figure 6B**). Its composition was most similar to that of the Built Environment (BLD) cluster, which consists almost exclusively of samples from anthropogenic surfaces (**Figure 2A**). This finding is consistent with previous reports indicating that urban microbiomes closely resemble human skin microbiomes[3].

#### Clusters consist of heterogeneous samples

Although most animal gut clusters exhibited high host specificity, the ANI cluster notably comprised gut samples from diverse hosts, including *Desmodus*, *Gymnogyps*, and *Ailuropoda* (**Figure 2D**). Considering that these animals are evolutionarily distant, inhabit distinct environments, and have markedly different dietary habits, the ANI cluster likely does not represent a specific animal or dietary pattern. Similarly, although approximately 70% of the samples in the HET cluster originated from anthropogenic environments, such as built-environment surfaces and wastewater, it also included samples with unclear commonalities, such as terrestrial samples and animal intestine microbiomes. It is possible that habitat types with limited sampling, lower sequencing quality, or particularly sparse microbiomes did not provide sufficient variability to robustly differentiate each habitat cluster via UMAP and subsequent clustering. Nevertheless, the ANI cluster was analyzed downstream as representative of animal gut environments, while the HET cluster, consisting predominantly (70%) of anthropogenic samples, was analyzed as representative of anthropogenic environments.
