## Supplementary Figure for "Planetary microbiome structure and generalist-driven gene flow across disparate habitats"

### Supplementary Figures

#### Supplementary Figure 1 | Benchmarking of taxonomic profile-based clustering models

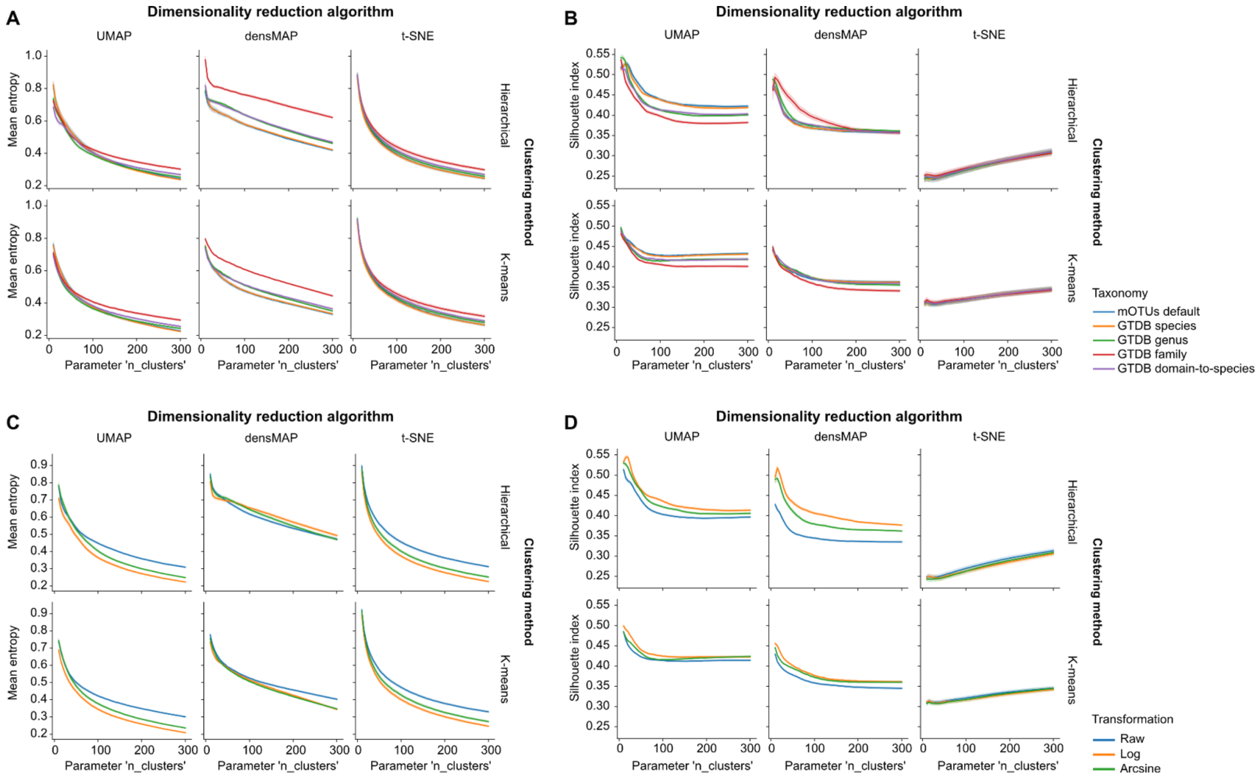


**(A–B)** Comparison of clustering models generated using different taxonomic resolutions. (A) Habitat entropy (lower scores represent better alignment with known broad habitat categories) and (B) Silhouette index (higher scores indicate better cluster separation) are compared across models based on species-level mOTUs, GTDB species, genus, family, and domain-to-species level profiles.

**(C–D)** Comparison of clustering models generated using different data transformations. (C) Habitat entropy and (D) Silhouette index are compared across models using raw, log-transformed, and arcsine-transformed data.

**Supplementary Figure 2 | Benchmarking of taxonomic, functional, and k-mer profile-based clustering models**

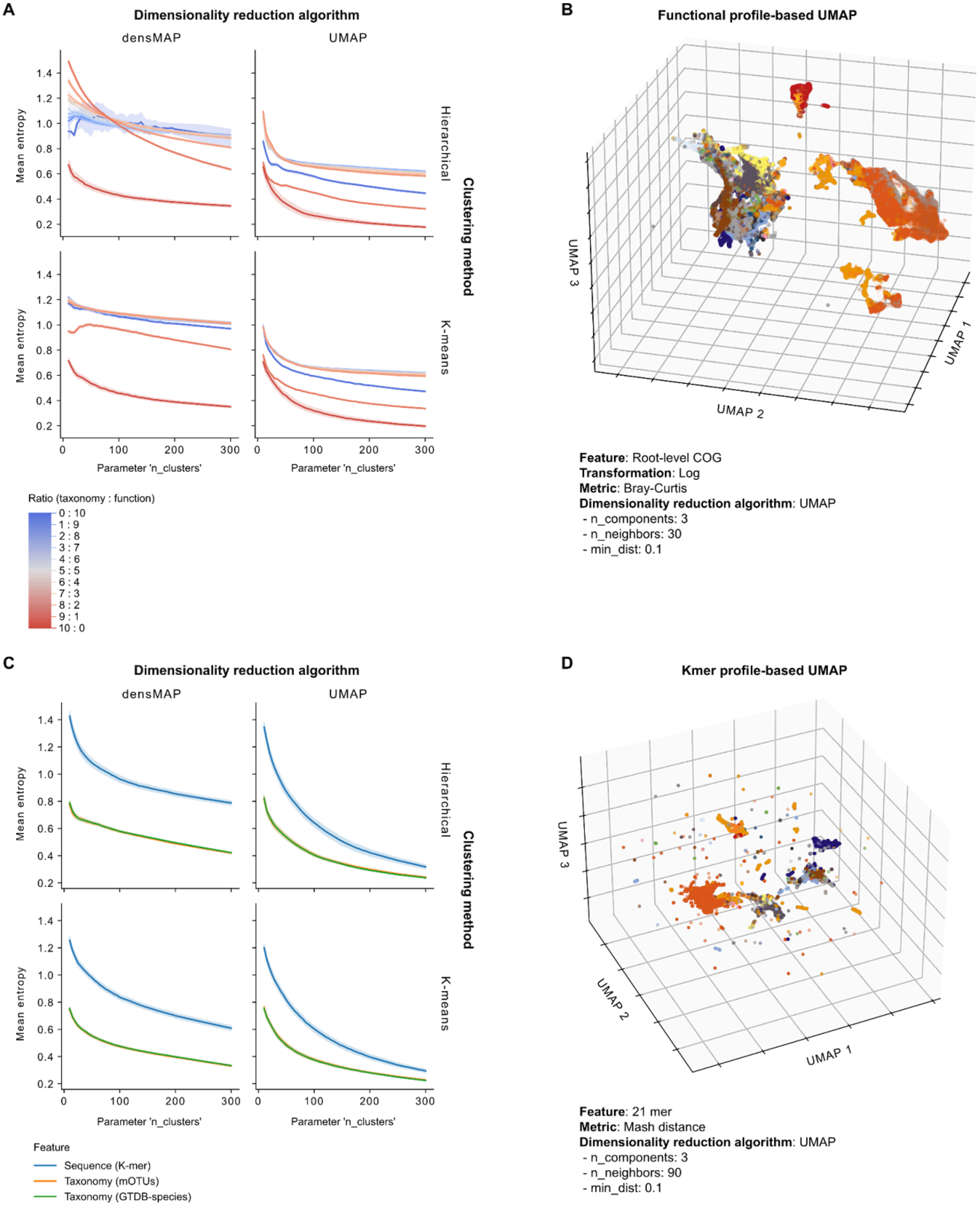


**(A)** Comparison of habitat entropy across clustering models based on functional profiles (blue), taxonomic profiles (red), and various ratios of combined functional-taxonomic profiles (blue-to-red gradient, **Methods**).

**(B)** 3D UMAP embedding of the clustering model built using the functional (root-level COG) profile.
**(C)** Comparison of habitat entropy for clustering models based on mash distance and taxonomic profiles.
**(D)** 3D UMAP embedding of the clustering model built using the mash distance profile. Datapoint colors in (B) and (D) represent sample microntology, matching the color scheme in Figure 1.

**Supplementary Figure 3 | Post-hoc refinement of the best-performing habitat clustering model**

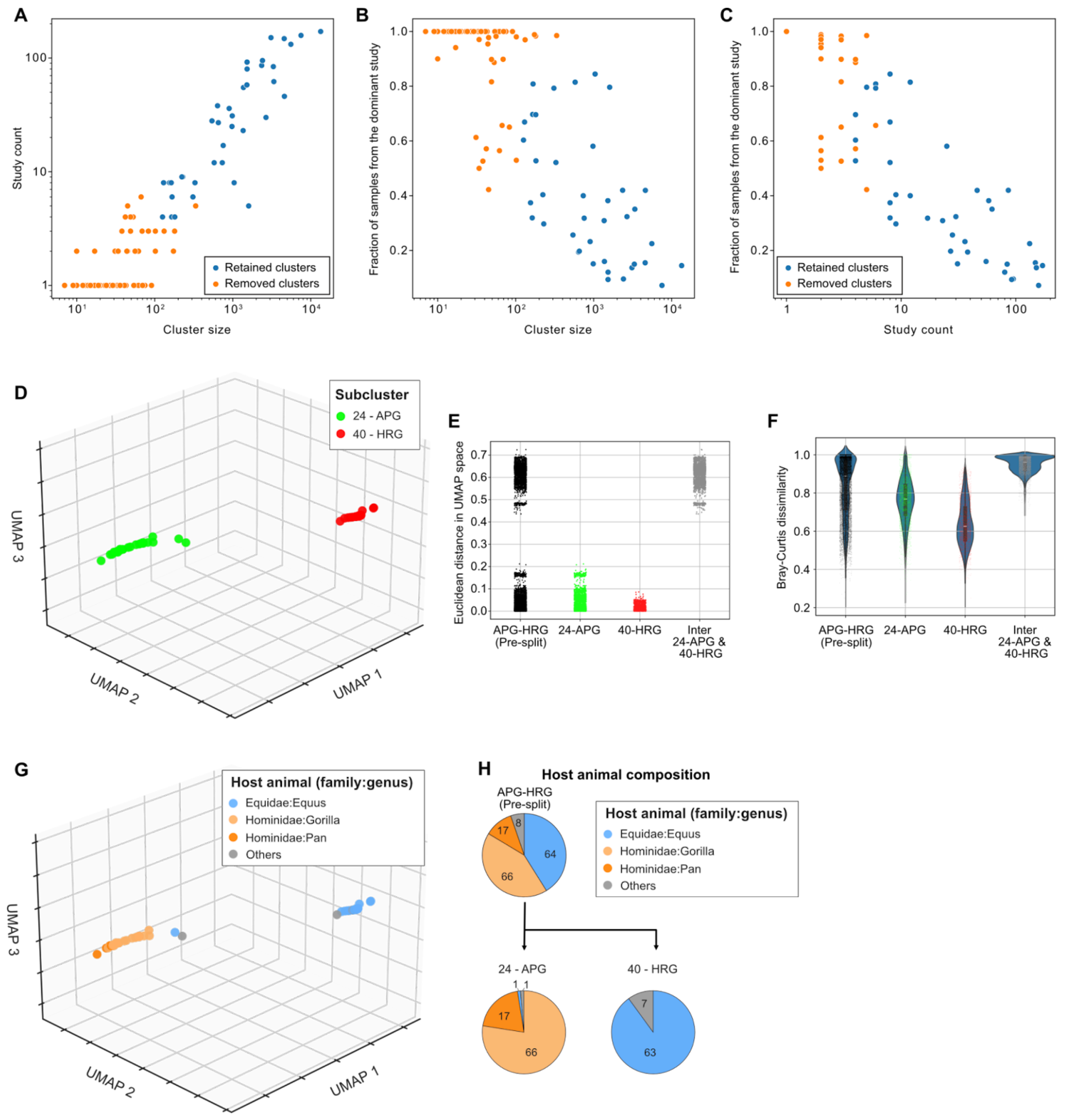


**(A-C)** Correlations of cluster size (number of samples within each cluster), study count (number of unique studies from which the samples were derived), and the fraction of samples from the dominant study for all clusters. The 39 retained clusters are highlighted as blue data points included in the final habitat clustering model.

**(D)** 3D UMAP embedding of the pre-split 24-APG and 40-HRG clusters.

**(E-F)** Comparison of inter- and intra-cluster distances for the pre- and post-split of the 24-APG and 40-HRG clusters. (E) Euclidean distances calculated in UMAP embedding, and (F) Bray-Curtis dissimilarity calculated from the original taxonomic profiles.

**(G)** UMAP embedding of 24-APG and 40-HRG clusters, visualized by host animal (family: genus).

**(H)** Host animal composition of 24-APG and 40-HRG clusters before and after the split.

#### Supplementary Figure 4 | UMAP embedding of the best-performing habitat cluster model

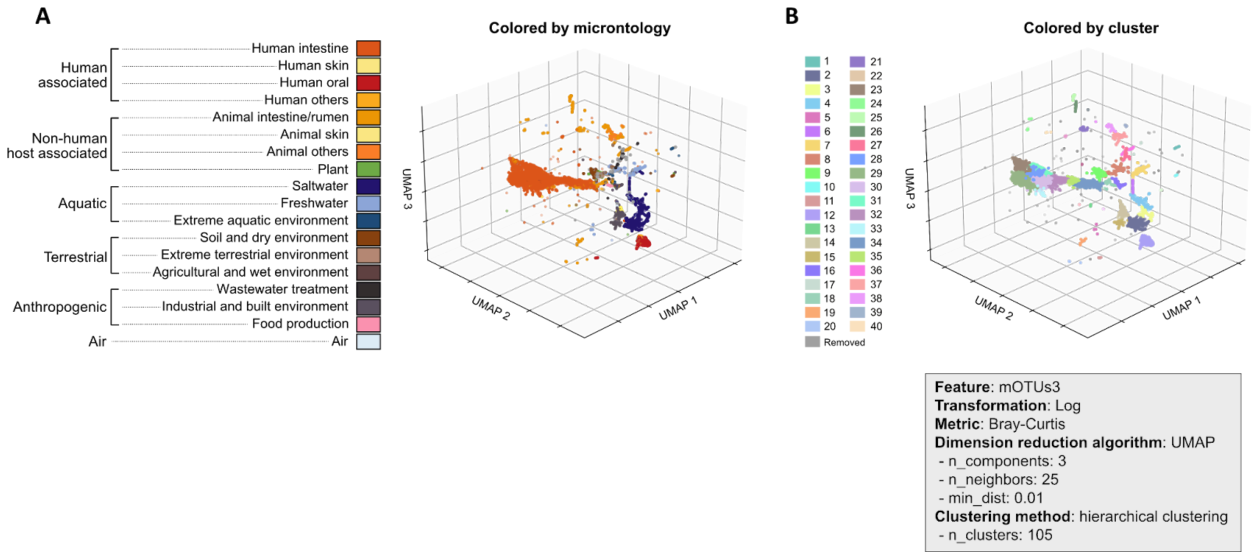


**(A-B)** 3D UMAP embedding of the best-performing habitat clustering model, colored by (A) microntology and (B) habitat cluster.

**Supplementary Figure 5 | Geographical and economic distribution of microbiome hosts within the human gut clusters.**

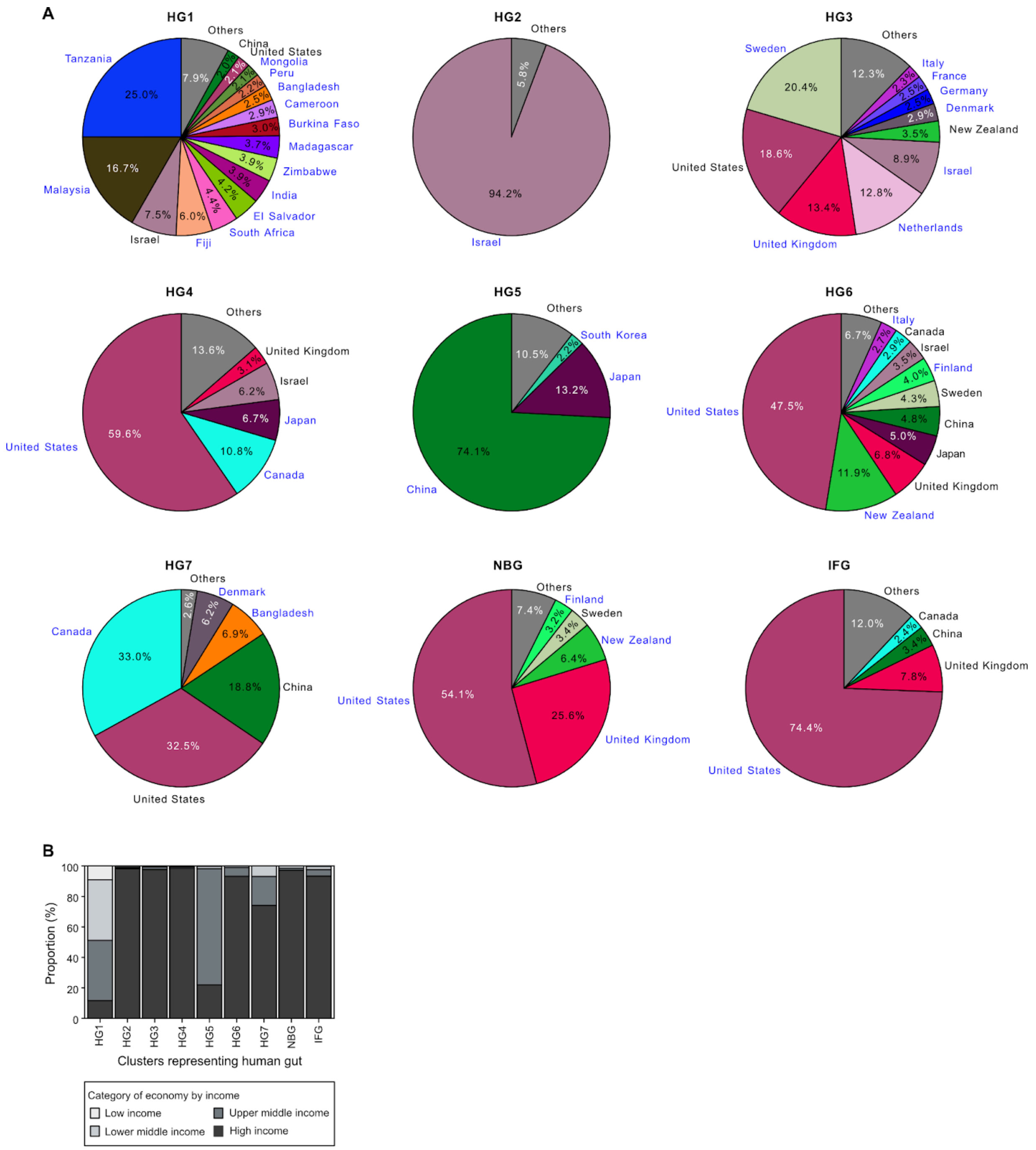


**(A)** Composition of countries where samples in each human gut cluster were collected. Blue text labels indicate countries that contributed a significantly higher proportion of samples to the respective cluster (Two-sided Fisher's test, Bonferroni-corrected $p < 0.1$, Odds ratio > 1). **(B)** Income categories of countries (The World Bank, 2024) by human gut clusters.

#### Supplementary Figure 6 | Habitat cluster model identifies host animals and putative sample contamination

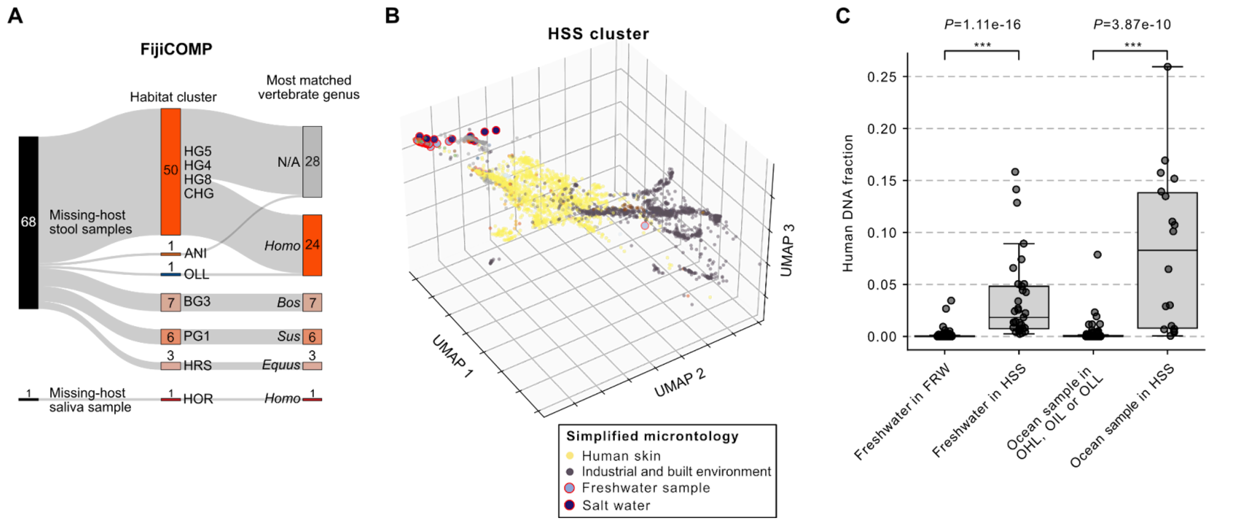


**(A)** Habitat classification and host DNA contamination analysis of samples lacking host annotations from the FijiCOMP dataset. These stool samples were mapped to habitat clusters, indicating their best matching vertebrate genus based on host DNA contamination in the Sankey diagram.

**(B)** UMAP embedding of the Human skin and surface cluster (HSS). Samples with metadata indicating freshwater or ocean environments are highlighted with red outlines.

**(C)** Comparison of human DNA content between freshwater and ocean samples classified into HSS and those assigned to clusters matching their habitat metadata (FRW, OIL, OLL, or OHL).

#### Supplementary Figure 7 | Phenotypic trait profiles at the habitat cluster level

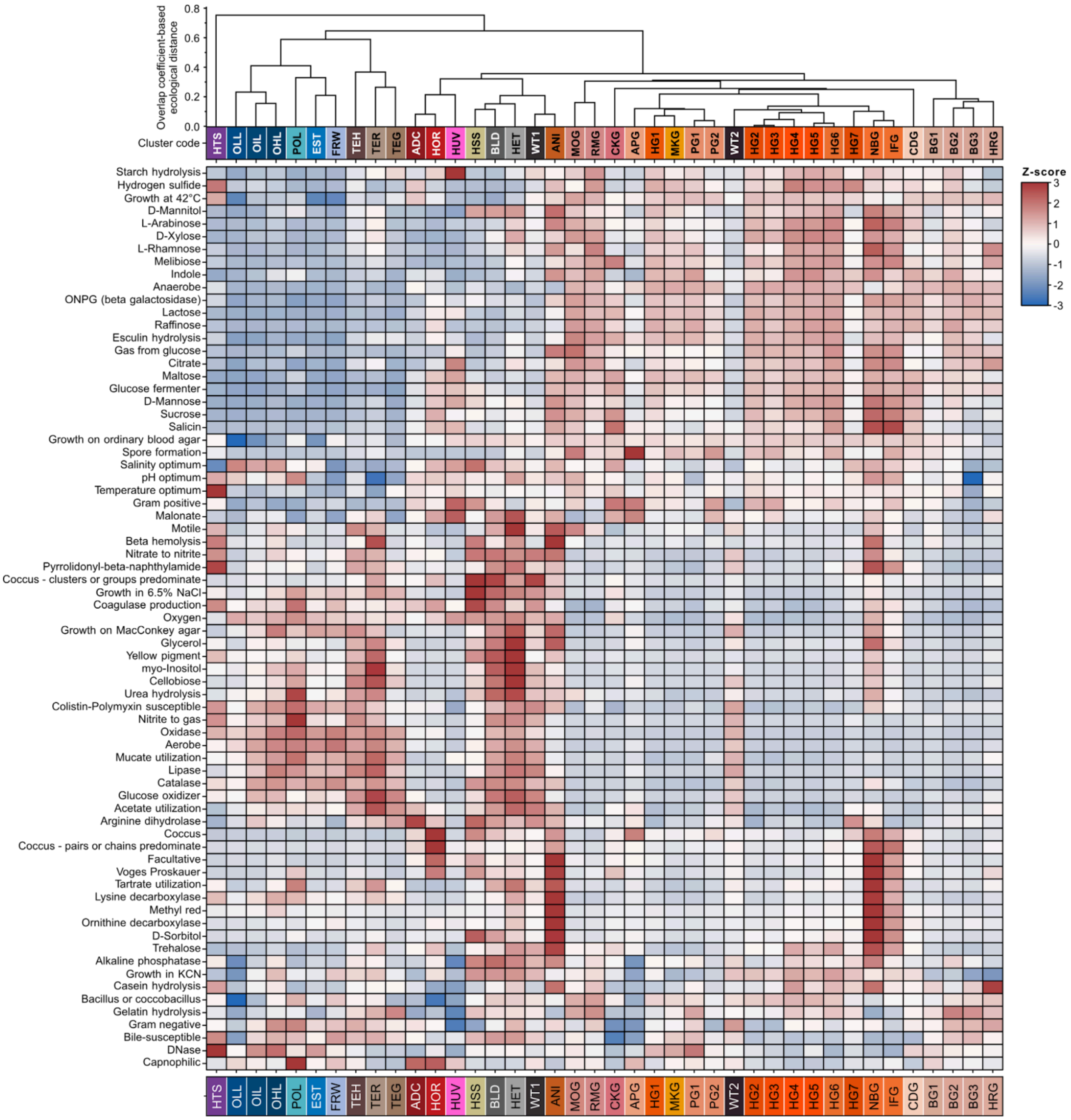


Heatmap illustrating the predicted phenotypic traits of species residing within habitat clusters. Genome-based predictions of microbial phenotypic traits were quantified at the habitat level by multiplying each species' trait prevalence with the habitat-specific mean relative abundance.

#### Supplementary Figure 8 | Rarefaction analysis of species and genus-level richness

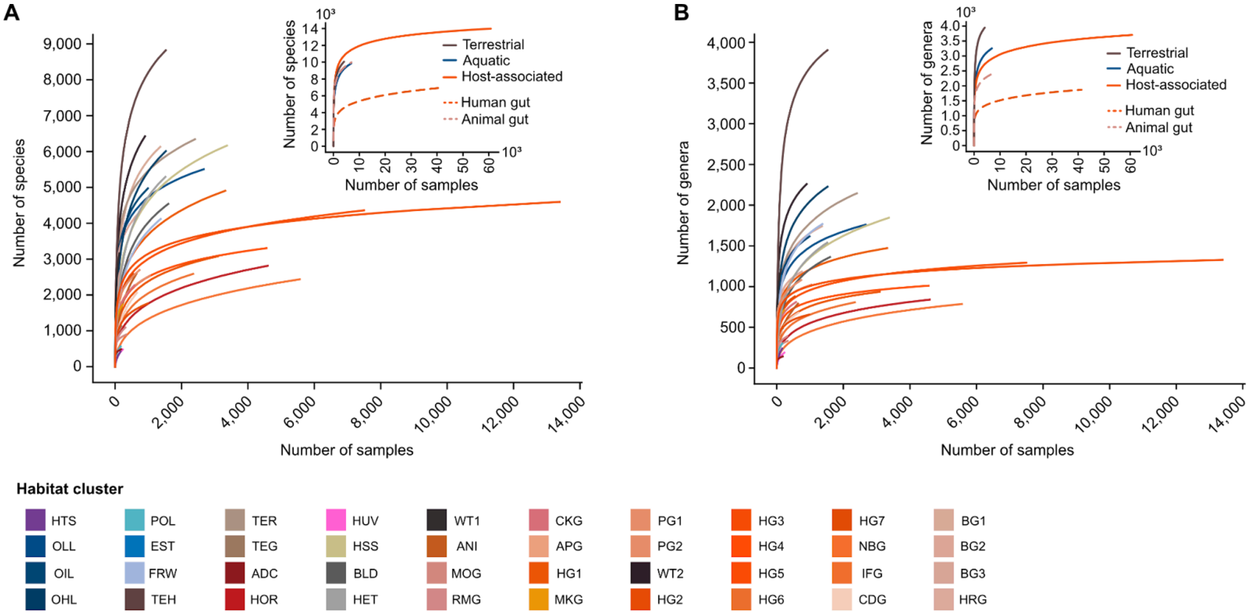


**(A-B)** Assessment of species (A) and genus-level (B) richness across simulated sampling efforts (rarefaction analysis). Insets show rarefaction curves after aggregating habitat clusters into terrestrial (**Figure 2A**-clade c), aquatic (**Figure 2A**-clade b), host-associated (**Figure 2A**-clade m), human gut (HG1–7, IFG, and NBG) and animal gut (ANI, MOG, RMG, CKG, APG, MKG, PG1–2, CDG, BG1–3, and HRG) categories. Human gut and animal gut are part of the host-associated category but are shown separately due to their relevance in microbiome research.

#### Supplementary Figure 9 | Schema of the generalism score calculation

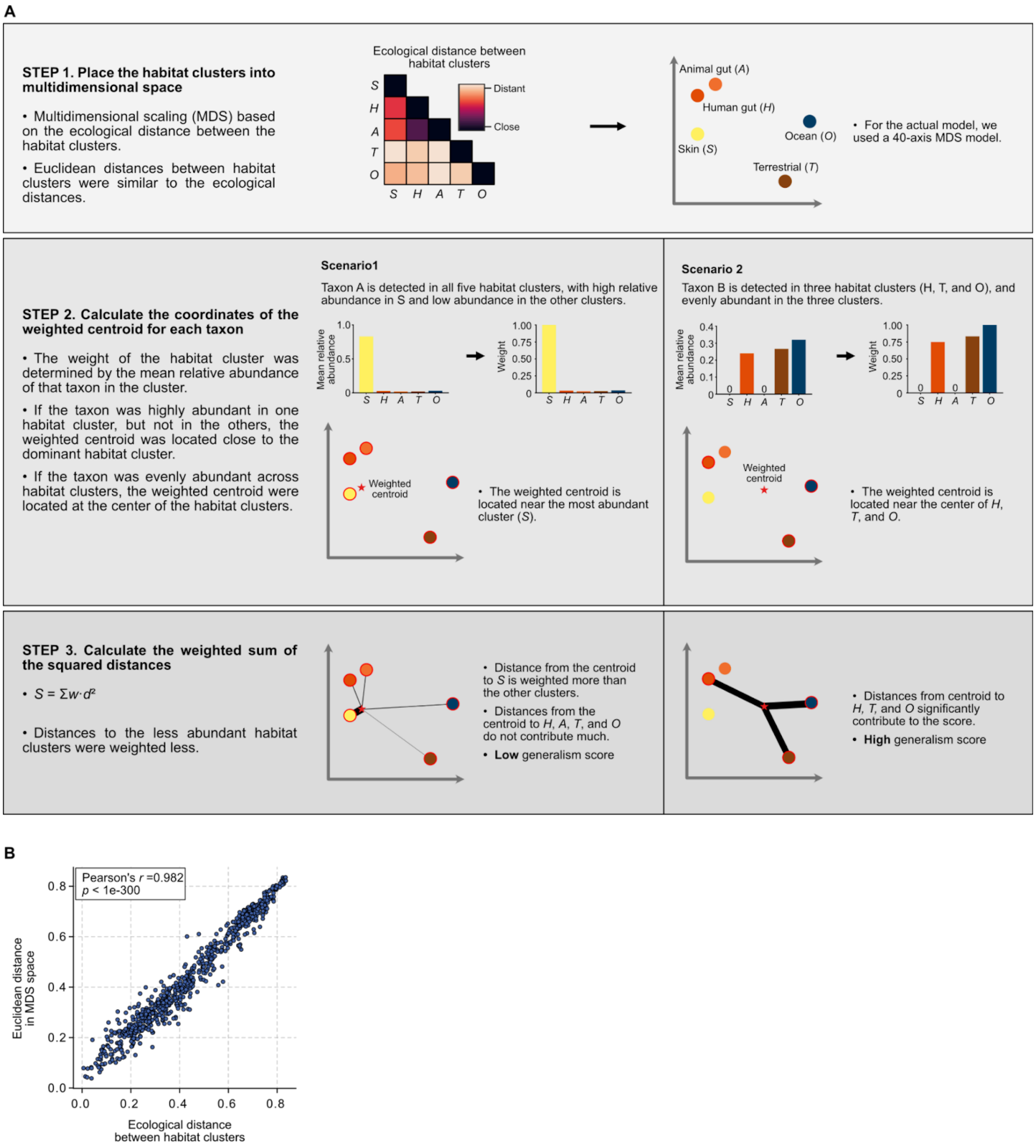


**(A)** Schema of the centroid-based generalism score calculation method, which considers the ecological distance between a microbe's habitat(s) and how evenly the microbe is distributed among them.

**(B)** The correlation between the overlap coefficient based phylogenetic distance and the Euclidean distance in MDS space was evaluated using Pearson's correlation coefficients.

**Supplementary Figure 10 | Genomic and phenotypic traits, and pathways associated with the generalism score**

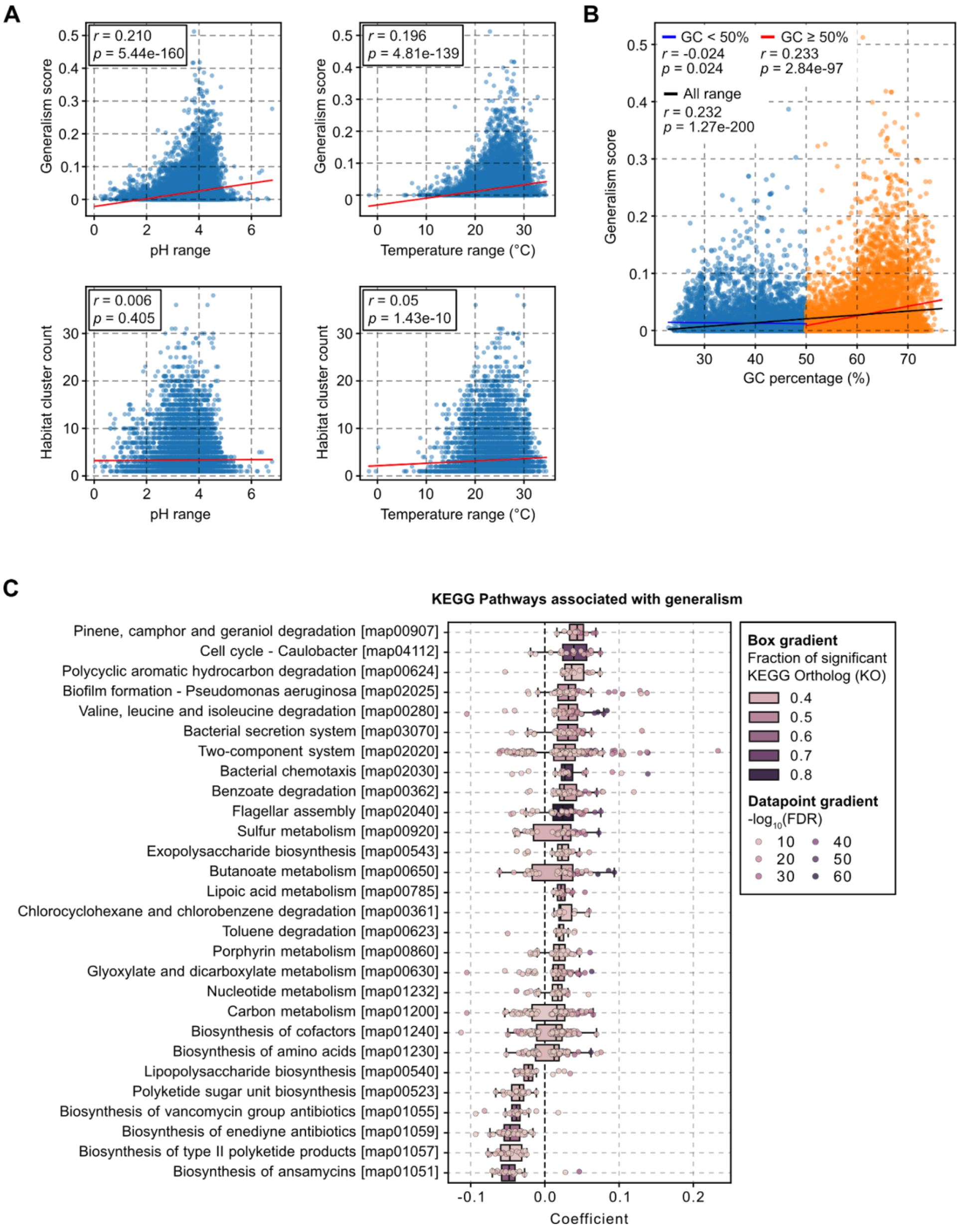


**(A)** Association of the generalism score (top panels) and the habitat cluster count (bottom panels) with the predicted pH range (left panels) and temperature range (right panels) are shown. Red lines indicate linear regression models. Pearson's correlation coefficients (*r*) and *p*-values are displayed.

**(B)** Association between GC content and the generalism score. Blue points and their regression line represent species with GC contents < 50%, whereas orange points and their regression line represent species with GC contents ≥ 50%. The black regression line indicates the correlation across all species. Pearson's correlation coefficients (*r*) and *p*-values are displayed.

**(C)** KEGG pathways (KPs) associated with prokaryotic generalism. Data points represent KEGG orthologs (KOs) within each pathway. Only KPs with more than 40% of their KOs significantly (FDR < 1e-10) enriched in either generalist or specialist species are shown.

#### Supplementary Figure 11 | Generalist involvement in HGT across ecological distances

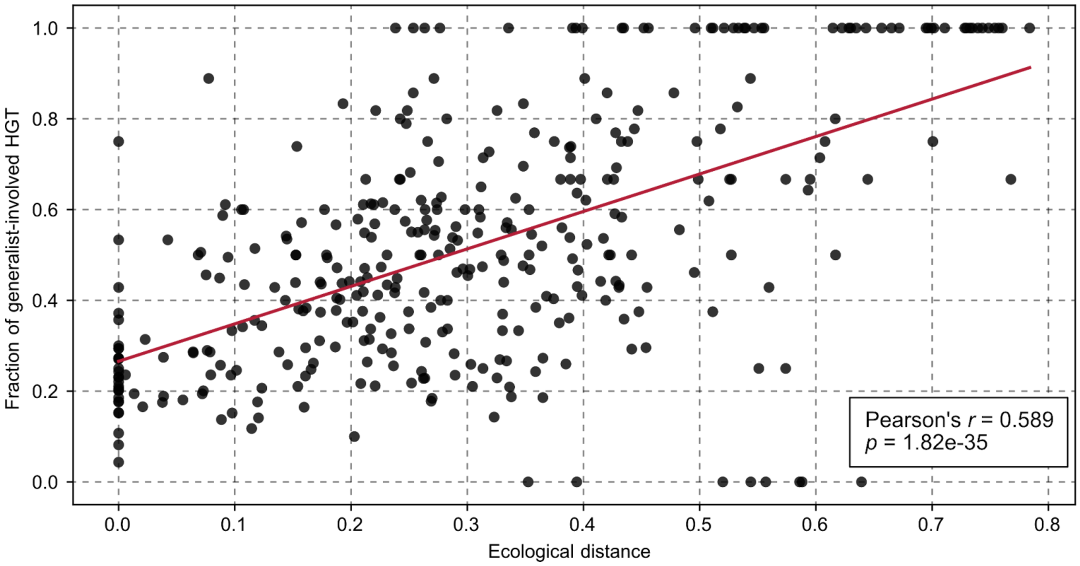


Scatter plot depicts the linear relationship (Pearson's *r* = 0.589, *p* = 1.82e-35) of ecological distance between habitat cluster pairs (> 1,000 MAGs with > 50% completeness) and their proportion of generalists (generalism score ≥ 0.1) involved in HGT.

#### Supplementary Figure 12 | Traits associated with abundance change in WT clusters

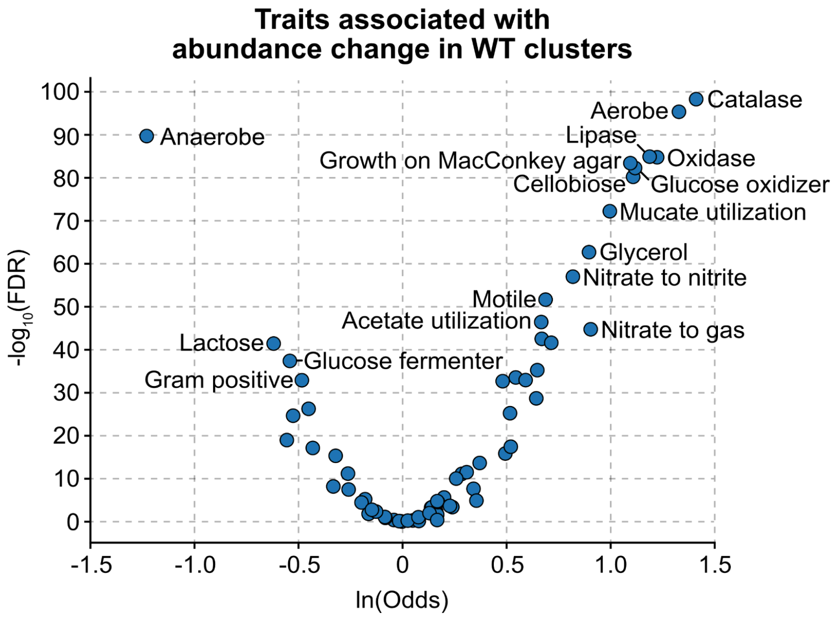


Volcano plot showing traits associated with abundance change, comparing gut to wastewater (WT) clusters. Positive odds indicate traits enriched in species with increased dominance in WT clusters.

**Supplementary Figure 13 | Horizontal gene transfer and antibiotic resistance**

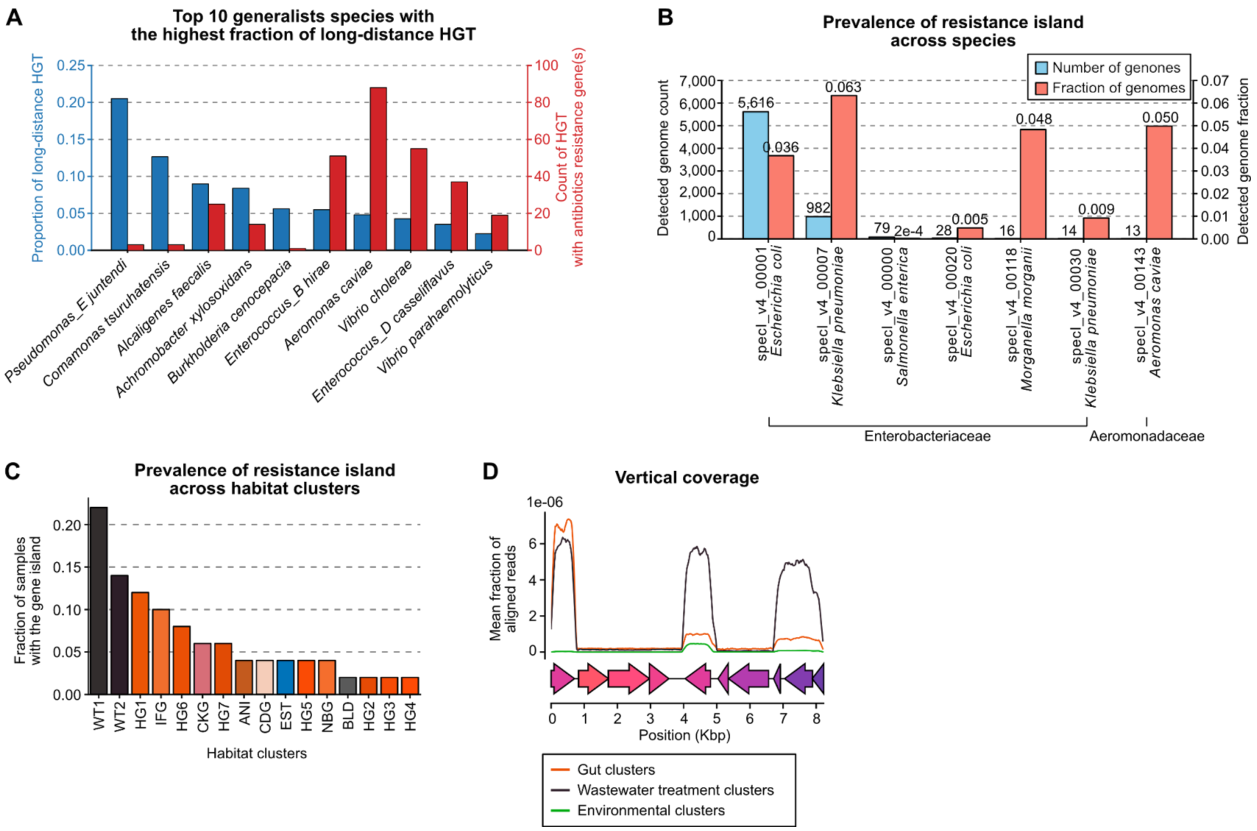


**(A)** For each of the 10 generalist species with the highest fraction of disparate-habitat HGT (between habitat cluster pairs with ecological distance > 0.5), the proportion of disparate-habitat HGT events and the number of ARG-containing HGTs are shown.
**(B)** Number and fraction of genomes containing the resistance island, organized by specI clusters. SpecI clusters with < 10 genomes containing the resistance island were excluded.
**(C)** Proportion of samples containing the resistance island (horizontal coverage > 90%) across habitat clusters.
**(D)** Average vertical coverage of the resistance island by metagenomic reads per environment.

#### Supplementary Figure 14 | Comparison between human ancient dental calculus and oral clusters

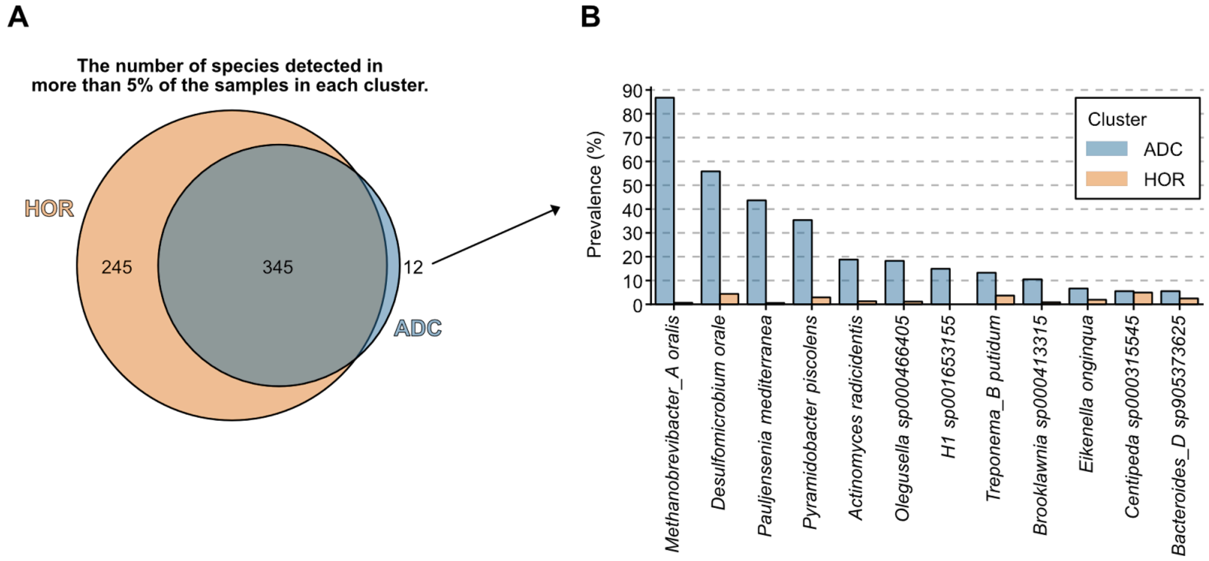


**(A)** Venn diagram representing the species overlap between HOR and ADC clusters, based on species that were detected in > 5% of the samples in each cluster.

**(B)** Prevalence of species that were detected with > 5% prevalence only in the ADC cluster.
